# Longitudinal Characterization of Cerebral Hemodynamics in the TgF344-AD Rat Model of Alzheimer’s Disease

**DOI:** 10.1101/2022.12.29.522215

**Authors:** Xing Fang, Chengyun Tang, Huawei Zhang, Jane J. Border, Yedan Liu, Seung Min Shin, Hongwei Yu, Richard J. Roman, Fan Fan

## Abstract

Alzheimer’s Disease (AD) is a global healthcare crisis. The TgF344-AD rat is an AD model exhibiting age-dependent AD pathological hallmarks. We confirmed that AD rats developed cognitive deficits at 6 months without alteration of any other major biophysical parameters. We longitudinally characterized cerebral hemodynamics in AD rats at 3, 4, 6, and 14 months. The myogenic responses of the cerebral arteries and arterioles were impaired at 4 months of age in the AD rats. Consistent with the *ex vivo* results, the AD rat exhibited poor autoregulation of surface and deep cortical cerebral blood flow two months preceding cognitive decline. The dysfunction of cerebral hemodynamics in AD is exacerbated with age associated with reduced cerebral perfusion. Further, abolished cell contractility contributes to cerebral hemodynamics imbalance in AD. This may be attributed to enhanced ROS production, reduced mitochondrial respiration and ATP production, and disrupted actin cytoskeleton in cerebral vascular contractile cells.

## INTRODUCTION

Alzheimer’s disease (AD) accounts for an estimated 60-80% of all cases of dementia ^1^. More than 6 million Americans over 65 years of age are living with AD in 2022. This disease cost the nation an estimated 321 billion dollars in Medicare costs and 271.6 billion dollars in unpaid care ^1^. The AD pathological hallmarks include abnormal accumulation of extraneuronal amyloid beta (Aβ) and intra-neuronal hyperphosphorylated tau resulting in neurodegeneration and cognitive decline. Emerging studies demonstrated that brain hypoperfusion is an early and persistent symptom of AD and AD-related dementias (ADRD) ^2-6^. Reduced perfusion in the neocortex and hippocampus has been reported to promote neuronal damage and dementia associated with poor cerebral blood flow (CBF) autoregulation, blood-brain barrier (BBB) leakage, and dysfunction of the neurovascular unit (NVU) in AD and aging, hypertension, and diabetes-related dementia ^7-14^. However, despite cerebral vascular dysfunction appearing earlier than Aβ and tau abnormalities, and much earlier than cognitive deficits, the relationship between Aβ accumulation and cerebral vascular dysfunction is unclear ^15,16^.

The TgF344-AD rat is an AD model that exhibits age-dependent increases in Aβ by overexpressing mutated human amyloid precursor protein (APP) with the Swedish mutation and human presenilin 1 (PS1) with the Δ exon 9 mutation onto the F344 genetic background ^17^. This AD model recapitulates the full spectrum of AD pathological hallmarks and starts to develop Aβ plaques, gliosis, and learning dysfunction as early as 6 months of age despite only containing mutant Aβ-producing genes ^17,18^. Other studies using this model demonstrated that the TgF344-AD rat also displayed BBB breakdown, neurofibrillary tangles, neuroinflammation, neurodegeneration, and diminished capability to increase CBF in response to hypercapnia ^17,19-21^. Dickie et al.^19^ reported that BBB water permeability increased with age from 13 to 21 months of age, but earlier in the TgF344-AD compared with F344 wildtype (WT) rats. Joo et al. ^20^ found that hypercapnia-induced increases in CBF were significantly reduced in cortical penetrating arterioles and venules in 9 months old TgF344-AD rats. Bazzigaluppi et al. ^22^ reported that 4 months old TgF344-AD rats exhibited a sustained reduction in CBF in association with vascular remodeling following L-NAME hypertension even though baseline CBF and vasodilator responses to hypercapnia were similar in AD and WT rats.

The brain has limited reserves of oxygen and glucose, although it is a highly energy-demanding organ ^23^. A recent study reported that glucose uptake in the entorhinal cortex and the hippocampus was significantly reduced in 9 months old TgF344-AD rats ^24^. Intact cerebral hemodynamics is critical to ensure brain perfusion and maintain normal neuronal function. Recent reports from the Alzheimer’s Disease Neuroimaging Initiative (ADNI), the Atherosclerosis Risk in Communities (ARIC), and the Genome-Wide Association Studies demonstrated that dementia in AD is associated with inactivating mutations in genes involved in CBF autoregulation ^25-28^. Poor CBF autoregulation is one of the essential vascular mechanisms resulting in brain hypoperfusion, neurodegeneration, and cognitive deficits ^4,29-31^. Autoregulation of CBF is modulated by the myogenic response (MR) of cerebral arteries and arterioles, which is an intrinsic property of vascular smooth muscle cells (VSMCs) that initiates vasoconstriction in response to elevations in pressure ^32-35^. The present study took advantage of this TgF344-AD rat model that contains Aβ-overproducing but not Tau-producing genes and longitudinally characterized cerebral hemodynamics at 3, 4, 6, and 14 months. We evaluated the MR of the freshly isolated middle cerebral artery (MCA) and parenchymal arteriole (PA) of AD and WT rats using a pressure myograph system. We compared autoregulation of both surface and deep cortical CBF in AD and WT rats using a laser-Doppler flowmeter (LDF), as well as cerebral perfusion using a laser speckle imaging system. Moreover, we also explored potential underlying mechanisms in primary VSMCs isolated from the MCAs of AD and WT rats by comparing cell contractile capability, production of reactive oxygen species (ROS), mitochondrial respiration and ATP production, and the actin-myosin contractile units.

## MATERIALS AND METHODS

### Animals

The TgF344-AD (F344-Tg (Prp-APP, Prp-PS1)19) rats were purchased from the Rat Resource & Research Center at the University of Missouri (Columbia, MO), and WT F344 rats (F344/NHsd) from ENVIGO (Indianapolis, IN). The two strains were inbred and maintained at the University of Mississippi Medical Center (UMMC). The present study was performed using male TgF344-AD and age-matched F344 rats at 3, 4, 6, and 14 months of age. The rats were provided food and water *ad libitum* and maintained in a centralized facility at the UMMC with a typical 12-hour light-dark cycle. All protocols in this study were approved by the Institutional Animal Care and Use Committee at the UMMC in accordance with the American Association for the Accreditation of Laboratory Animal Care, following the NIH guidelines.

### Baseline Biophysical Parameters

Baseline biophysical parameters, including body weight, plasma glucose, glycosylated hemoglobin A1c (HbA1c), and mean arterial pressure (MAP), were evaluated in 3, 4, 6, and 14 months TgF344-AD and WT rats following protocols we previously described ^9,36-39^. Briefly, rats were weighed, and blood samples were collected from the lateral tail vein in conscious rats 3 hours after the light-on cycle. Plasma glucose and HbA1c were examined using a Contour Next glucometer (Ascensia Diabetes Care, Parsippany, NJ) and an A1CNow^+^ system (PTS Diagnostics, Whitestown, IN), respectively. MAP was detected by implanting a telemetry transmitter (HD-S10, Data Sciences International, St. Paul, MN) into the femoral artery.

### Eight-arm Water Maze

All animals were validated by genotyping following protocols established by Cohen et al. ^17^ Longitudinal cognitive functions in separate groups of 3, 4, 6, and 14 months old male AD and WT rats were examined using an eight-arm water maze, as previously described ^9,37,40-42^, to validate hippocampal-based spatial learning and memory. This is a practical neurobehavioral test that takes advantage of the motivation of the subjects to escape without food deprivation ^43-46^. Briefly, the water maze was filled with tap water, maintained at 26°C ^47^. The animals were habituated to the test room for 1 hour in their home cages before the training phase. The training phase was performed on the first day, and the rats were trained to recognize and memorize the escape platform in one of the eight arms of the water maze. Eight trials were evaluated 2 and 24 hrs after the training phase. Data were recorded as the time to reach the platform (escape time) and errors (both the front and rear paws entering the wrong arm).

### Myogenic Response

The MRs of the MCA and PA of 3, 4, 6, and 14 months old male AD and WT rats were longitudinally evaluated. The MCAs and PAs were freshly isolated, as we previously described ^40,48^. Briefly, the rats were weighed and euthanized with 4% isoflurane and followed by rapid decapitation. The M2 segments of the MCA without any branches and lenticulostriate arterioles in the MCA territory that penetrate the brain parenchyma were carefully dissected and placed in ice-cold calcium-free physiological salt solution (PSS_0Ca_) supplemented with 1% bovine serum albumin (BSA) ^48-51^. Intact MCAs and PAs were then mounted onto glass cannulas in a pressure myograph chamber (Living System Instrumentation, Burlington, VT) that was filled with warmed (37 °C) and oxygenated (21% O_2_, 5% CO_2_, 74% N_2_) PSS solution containing calcium (PSS_ca_) ^9,48,52^. The MCAs and PAs were pressurized to 40 mmHg and 10 mmHg, respectively, for 30 minutes to develop a spontaneous tone. The inner diameters (IDs) of the MCAs and PAs in response to transmural pressure from 40 to 180 mmHg and 10 to 60 mmHg, respectively, were recorded by a digital camera (MU 1000, AmScope) mounted on an IMT-2 inverted microscope (Olympus, Center Valley, PA) to access the pressure-diameter relationships.

### Autoregulation of the Surface and Deep Cortical CBF and Brain Perfusion

Autoregulation of the surface and deep cortical CBF was evaluated in 3, 4, 6, and 14 months old male AD and WT rats following our previously optimized protocols ^40,53^. Briefly, the rats were anesthetized with Inactin (50 mg/kg; *i*.*p*.) and ketamine (30 mg/kg; *i*.*m*.). This combination has minimal impact on the CBF autoregulation and maintains baseline MAP in the physiological range in conscious rats ^10,40,54^. The rats were placed on a servo-controlled heating pad. The tracheal was cannulated with polyethylene (PE-240) tubing and connected to a ventilator (SAR-830, CWE Inc.). The femoral vein and artery were cannulated for drug delivery and MAP measurements, respectively. The head was secured in a stereotaxic device (Stoelting, Wood Dale, IL). The respiratory rate was adjusted to maintain an end-tidal P_CO2_ at 35 mmHg with a CO_2_ Analyzer (CAPSTAR-100, CWE Inc., Ardmore, PA). A closed translucent cranial window was created using a low-speed air drill, and a fiber-optic probe (91-00124, Perimed Inc., Las Vegas, NV) coupled to an LDF device (PF5010, Perimed Inc.) was placed above the cranial window for recording the surface cortical regional CBF. Another fiber-optic probe was implanted into the brain via a 1 mm hole to a depth of 1.5 - 2 mm for recording the deep cortical regional CBF ^9,37,55^. This hole was then sealed using bone wax. Baseline surface and deep cortical CBF were recorded at 100 mmHg. MAP was then elevated in steps of 20 mmHg to 180 mmHg by infusing phenylephrine (Sigma-Aldrich, St. Louis, MO). Then, the infusion of phenylephrine was withdrawn to return MAP to baseline. MAP was reduced in steps of 20 mmHg to 40 mmHg by graded hemorrhage.

Relative changes in CBF or brain perfusion in 3 and 14 months old male AD and WT rats were determined by a laser speckle imaging (LSI) system (RWD Life Science Co., Ltd, Shenzhen, China) as previously described ^56^. The surface CBF perfusion maps were captured at 60, 100, and 160 mmHg. Five regions of interest (ROI) were selected on each image, and the average LSI intensity normalized with the value at 100 mmHg was reported. All images were obtained under the same settings.

### Cell Contraction Assay

These experiments were performed using primary cerebral VSMCs isolated from the MCAs of male AD and WT rats following a protocol described earlier ^9,40,48,57^. Briefly, the MCAs were incubated with papain (22.5 U/mL) and dithiothreitol (2 mg/mL) at 37 °C for 15 minutes in the Tyrode’s solution (containing 145 mM NaCl, 6 mM KCl, 1 mM MgCl_2_.6H_2_O, 50 μM CaCl_2_.2H_2_O, 10 mM HEPES sodium salt, 4.2 mM NaHCO_3_, 10 mM Glucose; pH 7.4). After centrifugation at 1,000 rpm, the pellets of the digested vessels were resuspended in Tyrode’s solution and further digested with elastase (2.4 U/mL), collagenase (250 U/mL), and trypsin inhibitor (10,000 U/mL). The VSMCs were resuspended in the Dulbecco’s Modified Eagle’s Medium (DMEM, Thermo Scientific, Waltham, MA) containing 20% fetal bovine serum (FBS) and 1% penicillin/streptomycin after centrifugation at 1,500 rpm for 10 minutes at 37 °C. Cerebral VSMCs were isolated from 3-4 rats per strain.

Cell contractile capabilities were compared using primary cerebral VSMCs (passages 2 – 4) with a collagen gel-based cell contraction assay kit (CBA-201, Cell Biolabs, San Diego, CA) following our optimized protocol ^9,40,48,53,57,58^. Briefly, cells (3□×□10^6^ cells/mL) were seeded into a 24-well plate pre-coated with Cell-Tak Cell and Tissue Adhesive (354242, 3.5 μg/cm^2^; Corning Inc. Corning, NY). One group of cerebral VSMCs isolated from WT rats was pretreated with amyloid-beta peptides 1-42 (Aβ (1-42); 0.1 μM and 1 μM in FBS- and antibiotics-free culture medium; ANASPEC, Fremont, CA). The cell contraction matrix was prepared by mixing cell suspension and collagen gel working solution on the ice at a 1:4 ratio. After collagen polymerization, the cell-gel mixture (0.5 mL, containing 3 × 10^5^ cells) was supplied with DMEM (1 mL) containing vehicle, 0.1 μM or 1 μM Aβ (1-42), respectively, and incubated for 24 hours to develop cell contractile force. The stressed matrix was then gently released using a sterile spatula (time 0), followed by adding 1 mL of DMEM containing vehicle, 0.1 μM or 1 μM Aβ (1-42), respectively, and stimulated for 2 hours (time 120) to initiate the cell contraction. Changes in the size of the detached collagen gel (contraction index) were imaged at time 0 and time 120 and quantified with PowerPoint 2016 (Microsoft Corporation, Redmond, WA). Experiments using primary cerebral VSMCs were performed in triplicate and repeated three times.

### ROS Production

These experiments were performed using primary cerebral VSMCs (passages 2 – 4) isolated from the MCAs of male AD and WT rats. ROS production was detected with dihydroethidium (DHE; D11347, Thermo Scientific), and mitochondrial ROS production was measured using the MitoSOX™ Red Mitochondrial Superoxide Indicator kit (M36008; Thermo Fisher Scientific), as previously described ^57,58^. Briefly, cerebral VSMCs were cultured on a 6-well plate, washed with phosphate-buffered saline (PBS) after being firmly attached, and incubated with DHE (10 μM) or MitoSOX™ (5 μM) at room temperature for 10 min, respectively. Cells incubated with MitoSOX™ were then fixed with 3.7% paraformaldehyde and applied with an anti-fading mounting medium with 4′,6-diamidino-2-phenylindole (DAPI; H-1200, Vector Laboratories, Burlingame, CA). Live or fixed cell images were captured at an excitation/emission (in nm) of 405/590 and 396/610, respectively ^57,59^ using the Lionheart automated live-cell imager (BioTek Instruments, Inc., Winooski, VT) and a fluorescence microscope (Nikon, Melville, NY), respectively. Quantitation of relative ROS or mitochondrial ROS production was determined with Image J software (https://imagej.nih.gov/ij/download.html) by comparing the mean intensities of the red fluorescence ^57,58^. Experiments using primary cerebral VSMCs were performed in triplicate and repeated three times.

### Mitochondrial Respiration and ATP production

These experiments were performed using primary cerebral VSMCs (passages 2 – 4) isolated from the MCAs of male AD and WT rats. Mitochondrial respiration and ATP production were evaluated using the Seahorse XF^e^24 Extracellular Flux Analyzer (Agilent, Santa Clara, CA) by comparing the oxygen consumption rate (OCR) as we have previously described ^57,58^. Briefly, cerebral VSMCs (5 × 10^3^/well) were placed on an XF^e^24 plate pretreated with Corning Cell-Tak Cell and Tissue Adhesive. After incubation with XF base medium, OCR was measured before and after sequential administration of oligomycin (1 μM; Sigma-Aldrich), carbonyl cyanide-4-(trifluoromethoxy) phenylhydrazone (2 μM; FCCP, Sigma-Aldrich), rotenone (0.5 μM; Sigma-Aldrich) and antimycin-A (0.5 μM; Sigma-Aldrich). Results generated by the Agilent Seahorse XFe24 Analyzer Wave 2.4 software were normalized to protein concentrations in each well, as we described previously ^57,58^. Experiments using primary cerebral VSMCs were performed in triplicate and repeated three times.

### Immunocytochemistry

These experiments were performed using primary cerebral VSMCs (passages 2 – 4) isolated from the MCAs of male AD and WT rats. Cells were seeded onto a 4-well chamber slide, fixed with 3.7% paraformaldehyde, and permeabilized with 0.1% Triton-100 in PBS, as we previously described ^9,57,58^. After blocking with 1% BSA, the cells were incubated with Alexa Fluor 488 phalloidin (1:40; A12379, Thermo Fisher Scientific, Waltham, MA) to observe the shape and structure of F-actin. The cells were co-incubated with anti-myosin light chain (phospho S20) antibody (MLC; 1:300; ab2480, Abcam, Cambridge, MA) followed by Alexa Fluor 555 goat anti-rabbit (A-21428, Thermo Fisher Scientific). The slides were applied with an anti-fade mounting medium with DAPI (Vector Laboratories), coverslipped, and imaged using a Nikon C2 laser scanning confocal head mounted on an Eclipse Ti2 inverted microscope (Nikon, Melville, NY). MLC intensity and the MLC/F-actin ratios (of positively stained cells) were measured using the NIS-Elements Imaging Software 4.6 (Nikon), as we previously reported ^36,39^. Experiments using primary cerebral VSMCs were performed in triplicate and repeated three times.

### Statistical Analysis

Data are presented as mean values ± standard error of the mean (SEM). The statistical analysis was performed using GraphPad Prism 9 (GraphPad Software, Inc., La Jolla, CA). Significances between WT and AD rats in corresponding values in body weight, plasma glucose, HbA1c, MAP, eight-arm water maze, myogenic response, CBF autoregulation, and brain perfusion were compared with a two-way ANOVA for repeated measures followed by a Holm-Sidak *post hoc* test as described previously ^39,54,60^. Significance of the difference in corresponding values in cell contractility between cells isolated from WT (treated with vehicle and two doses of Aβ (1-42) and AD rats was compared with one-way ANOVA followed by a *post hoc* Tukey test. Significances in corresponding values of changes in ROS production, mitochondrial superoxide production, mitochondrial dynamic function, and immunocytochemistry between AD and WT were compared using an unpaired *t*-test. *P* < 0.05 was considered significant.

## RESULTS

### Baseline Biophysical Parameters

As presented in **Table 1**, body weights, plasma glucose, HbA1c, and MAP were similar between AD and age-matched WT rats across all ages, suggesting the cerebral vascular and cognitive dysfunction in the TgF344-AD rats were likely due to overexpression of Aβ by mutant APP and PS1 transgenes but not rather than diabetes or hypertension, consistent with previous observations ^17,22^.

**TABLE 1.** Baseline Biophysical Parameters. Comparison of body weights, plasma glucose and glycosylated hemoglobin (HbA1c) concentrations, and mean arterial pressure (MAP) in 3, 4, 6, and 14 months old wildtype (WT; F344) and TgF344-AD (AD) rats. Mean values ± SEM are presented. N = 4-12 rats per group. *P* < 0.05 was considered significant.

| Parameters | Age | WT | AD |
| --- | --- | --- | --- |
| **Body Weight (g)** | 3 m | 255.45 ± 7.28 | 243.50 ± 11.20 |
|  | 4 m | 296.75 ± 3.75 | 301.22 ± 13.13 |
|  | 6 m | 342.67 ± 8.26 | 351.33 ± 9.08 |
|  | 14 m | 435.92 ± 7.69 | 458.86 ± 12.89 |
| **Plasma Glucose (mg/dL)** | 3 m | 113.00 ± 3.11 | 116.00 ± 4.51 |
|  | 4 m | 113.50 ± 2.90 | 112.75 ± 2.43 |
|  | 6 m | 112.00 ± 3.83 | 113.25 ± 4.01 |
|  | 14 m | 112.00 ± 2.80 | 113.75 ± 4.03 |
| **HbA1c (%)** | 3 m | 4.25 ± 0.18 | 4.18 ± 0.03 |
|  | 4 m | 4.20 ± 0.09 | 4.38 ± 0.13 |
|  | 6 m | 4.15 ± 0.10 | 4.38 ± 0.14 |
|  | 14 m | 4.35 ± 0.09 | 4.25 ± 0.09 |
| **MAP (mmHg)** | 3 m | 109.00 ± 7.18 | 114.04 ± 5.53 |
|  | 4 m | 125.54 ± 3.74 | 115.51 ± 5.83 |
|  | 6 m | 121.33 ± 5.17 | 116.36 ± 9.00 |
|  | 14 m | 132.25 ± 4.09 | 134.00 ± 6.68 |

### Eight-arm Water Maze

We next examined spatial learning and memory phenotypes in 3, 4, 6, and 14 months old male WT and TgF344-AD rats using an eight-arm water maze. Consistent with previous reports ^17^, the AD rats started to display cognitive deficits at 6 months of age (**Fig.1)**. Six and 14 months old AD rats spent a longer time (**Fig.1C, 1D)** and had more errors (**Fig.1E)** than relative controls, indicating learning dysfunction and impaired short- and long-term memory.

**Figure 1.**
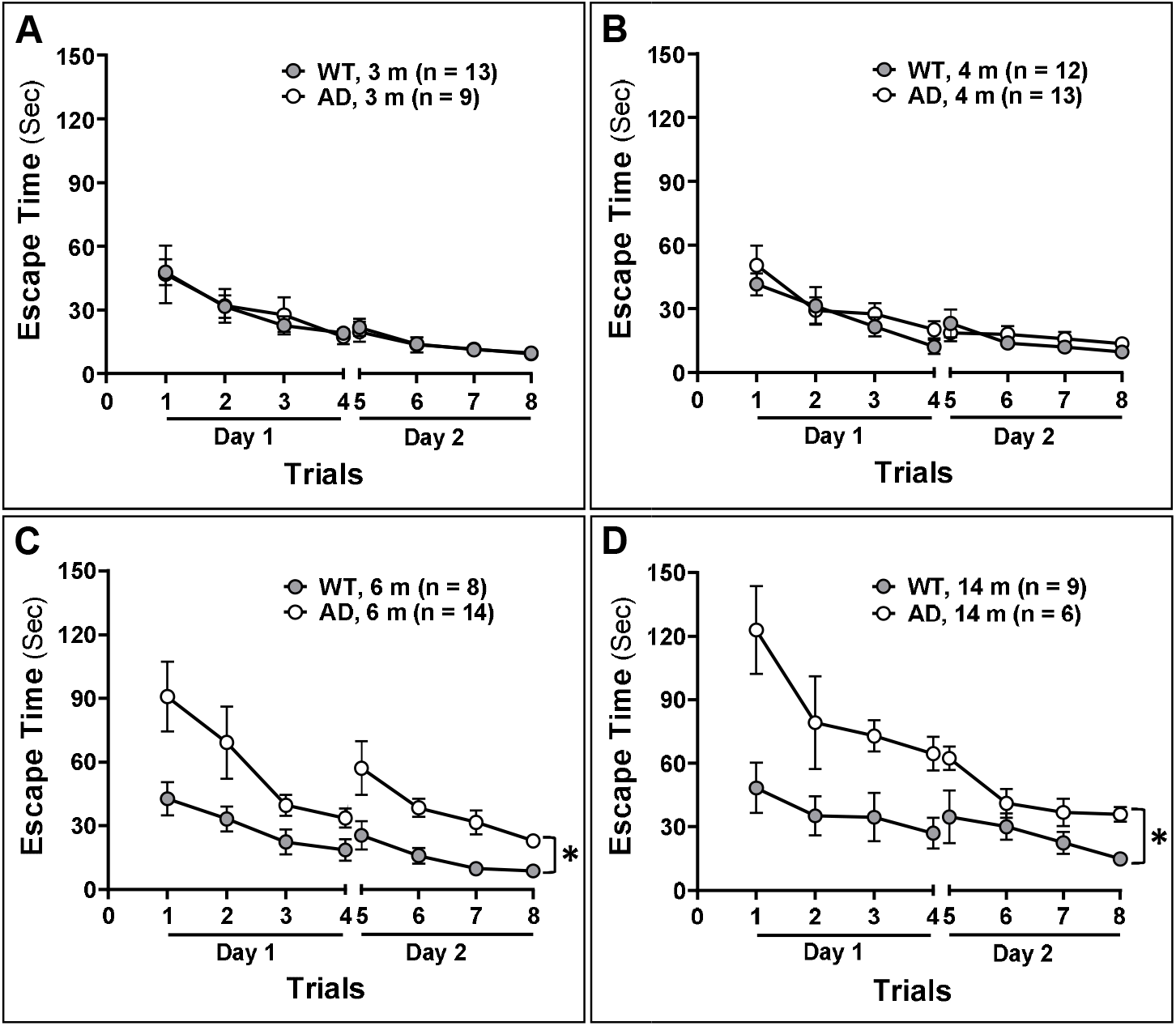
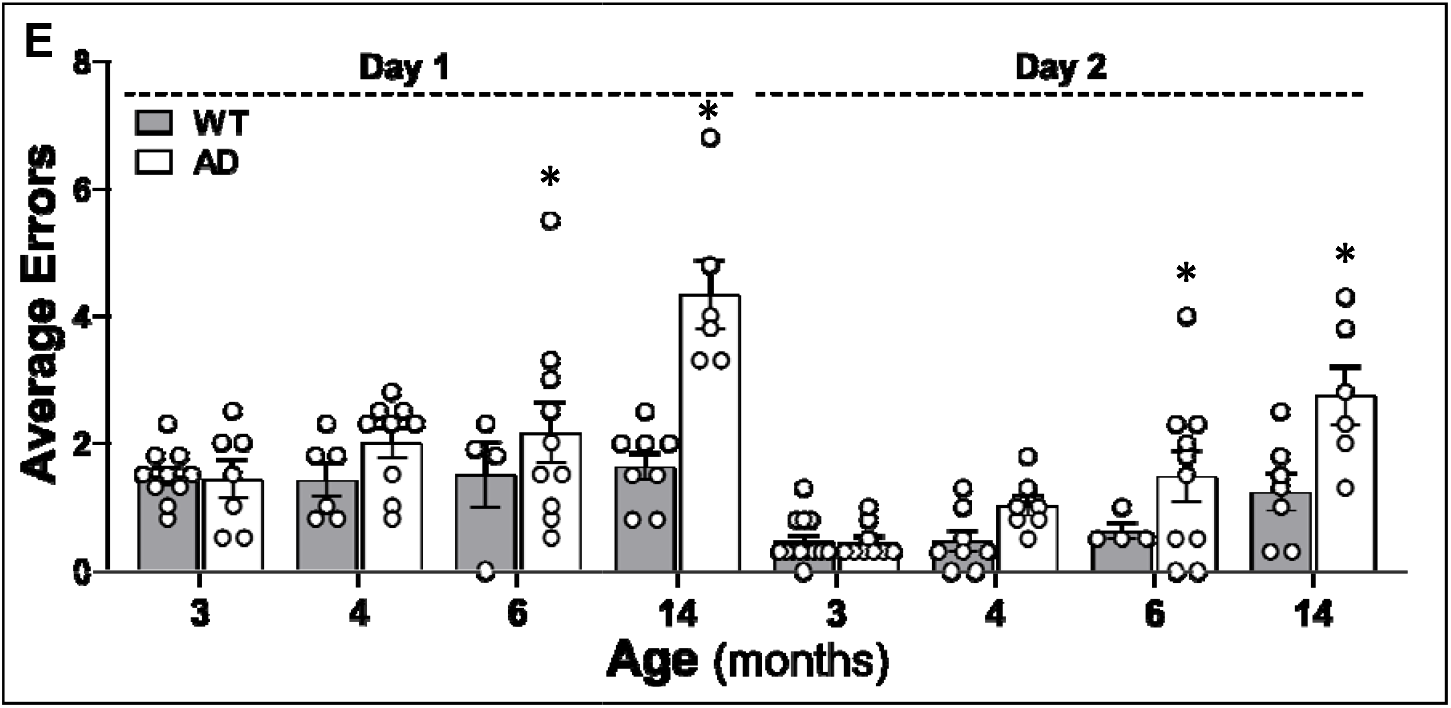
Validation of Cognitive Phenotype of the TgF344-AD (AD) rats. Hippocampal-based spatial learning and memory phenotype was evaluated in 3, 4, 6, and 14 months old wildtype (WT; F344) and AD rats. **A-D**. Time to reach the platform per trial (escape time) in a 2-day eight-arm water maze test. Numbers in parentheses indicate the number of rats studied per group. **E**. The total number of errors per trial. N = 6-14 rats per group. Mean values ± SEM are presented. An open circle represents a single value obtained from an individual animal. * indicates *P* < 0.05 from the corresponding values in age-matched WT rats.

### Myogenic Response

The MR of the MCA of 3, 4, 6, and 14 months old male AD and WT rats were longitudinally evaluated. As depicted in **Fig.2A-D**, The MCA of WT and AD rats exhibited a similar vasoconstrictive response to increases in perfusion pressure from 40-180 mmHg at 3 months of age (**Fig. 2A**). Interestingly, while the MR of the MCA of WT rats remained intact at 4, 6, and 14 months of age (**Fig. 2B-D**), AD rats started to lose the vasoconstrictive responses to pressures at 4 months, as MCA only constricted by 5.58 ± 2.67 % and 9.64 ± 2.87 % when transmural pressures were elevated from 40 to 100 and 140 mmHg, respectively, compared to 11.92 ± 1.66 % and 18.78 ± 3.05 % in WT when pressures were increased over the same ranges. The impaired MR was exacerbated with age in AD rats. AD vessels exhibited forced dilation over the pressure range of 120-180 mmHg at 4 and 6 months and 100-180 mmHg at 14 months of age.

**Figure 2.**
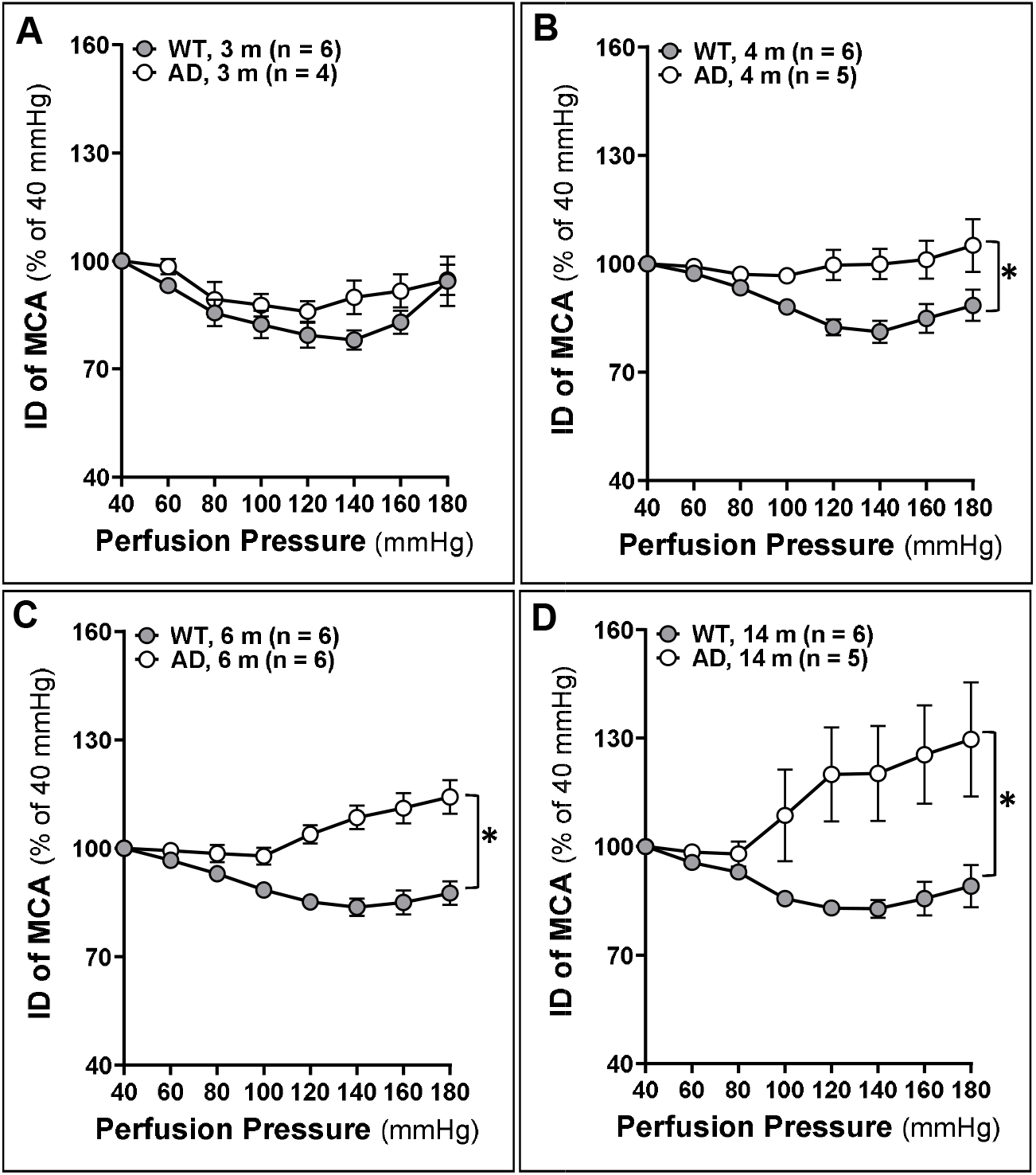
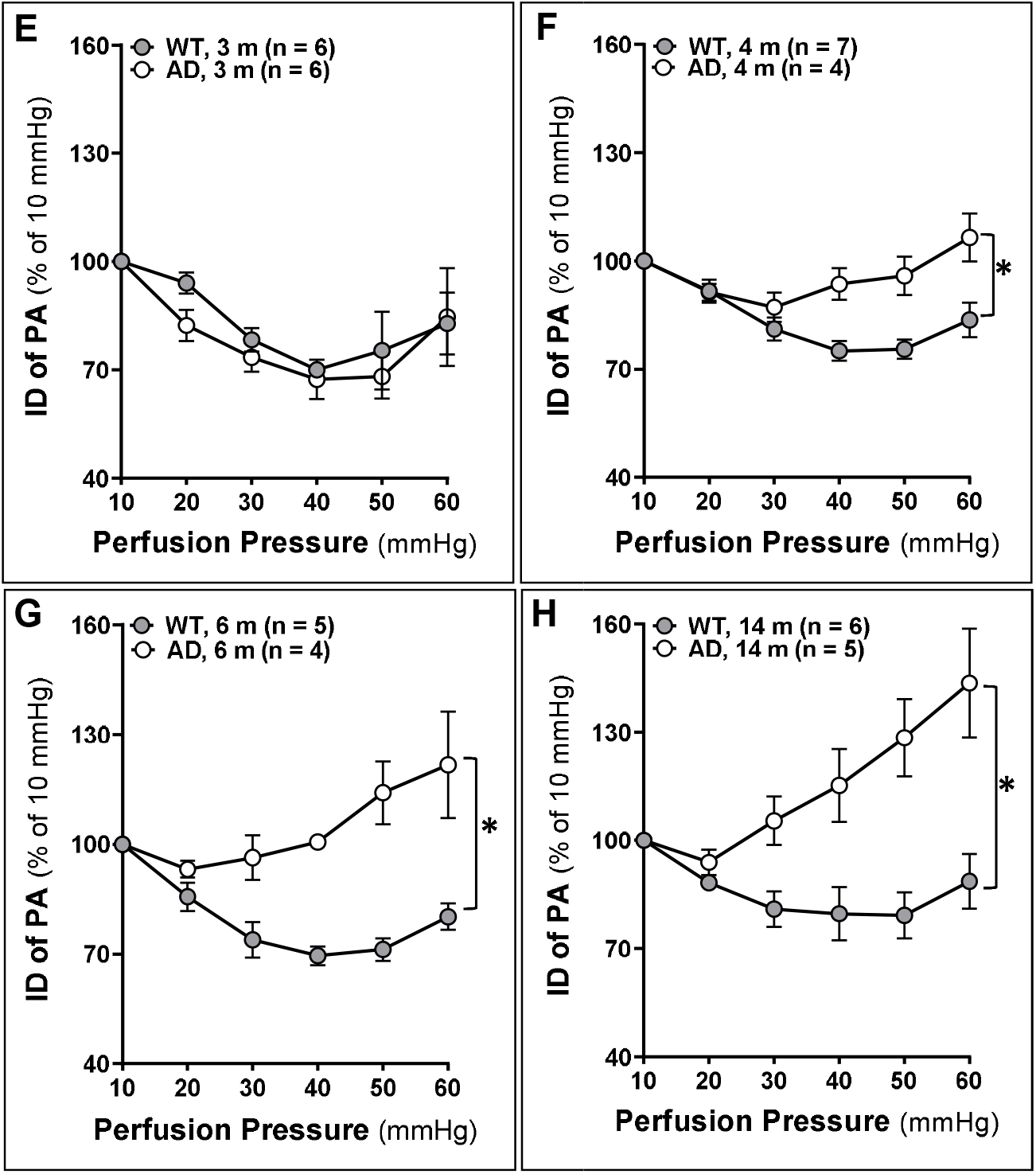
Myogenic Response. Comparison of the myogenic response of the middle cerebral artery (MCA) and parenchymal arteriole (PA) in 3, 4, 6, and 14 months old wildtype (WT; F344) and TgF344-AD (AD) rats. **A-D**. Comparison of the myogenic response of the MCA as of % constriction to 40 mmHg. **E-H**. Comparison of the myogenic response of the PA as of % constriction to 10 mmHg. Mean values ± SEM are presented. Numbers in parentheses indicate the number of rats studied per group. * indicates a significant difference (*P* < 0.05) from the corresponding values in age-matched WT rats.

The MR of the PA of 3, 4, 6, and 14 months old male AD and WT rats were also compared. Similar to what was seen in the MCA, PAs isolated from AD rats began to display an impaired MR at the age of 4 months and exacerbated with age (**Fig. 2E-H**). We found that the MR of the PA was similar in WT and AD rats at 3 months of age. However, impaired MR of PA was observed in 4 months old AD rats compared with WT rats. The impaired MR of PA was also exacerbated with age in AD rats. AD vessels exhibited forced dilation over a pressure range of 40-60 mmHg in 4 and 6 months old rats and 30-60 mmHg in 14 months old AD rats.

### CBF Autoregulation and Brain Perfusion

Surface cortical CBF autoregulation in 3, 4, 6, and 14 months male AD and WT rats was compared. As presented in **Fig. 3A-D**, WT and AD rats exhibited a similar intact surface cortical CBF autoregulation at 3 months of age. Interestingly, while the surface cortical CBF autoregulation in WT rats remained intact at 4, 6, and 14 months of age, AD rats failed to autoregulate beginning at 4 months of age, as CBF increased by 22.18 ± 1.39 % (WT) vs. 40.22 ± 5.63 % (AD) when perfusion pressure increased from 100 to 160 mmHg and decreased by 8.18 ± 1.95 % (WT) vs. 19.82 ± 1.31 % (AD) when perfusion pressure reduced from 100 to 60 mmHg, respectively. The impaired surface CBF autoregulation was exacerbated with age in AD rats. AD vessels exhibited autoregulatory breakthroughs above 140 and 120 mmHg at 4, and 6 or 14 months of age, respectively.

**Figure 3.**
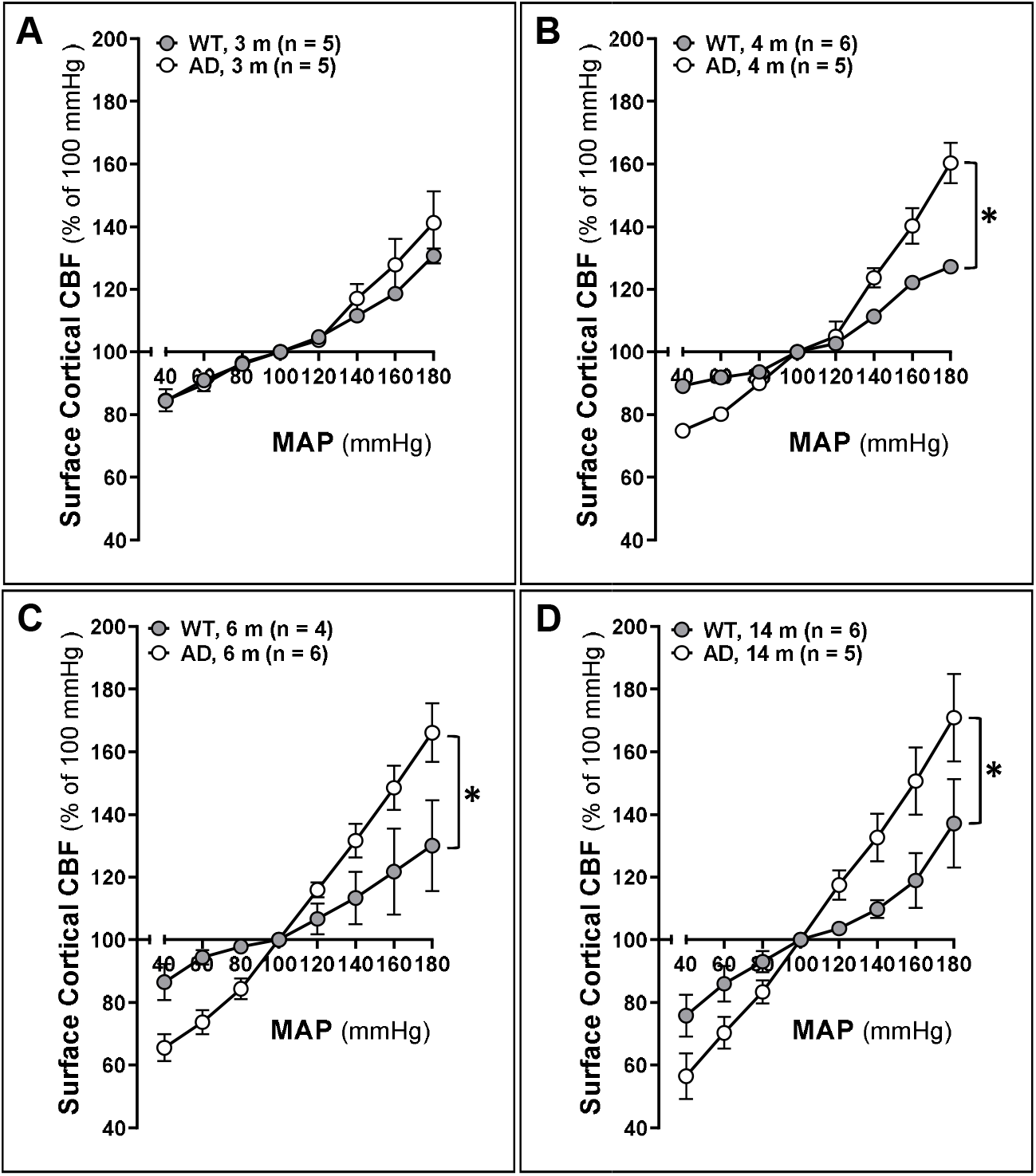
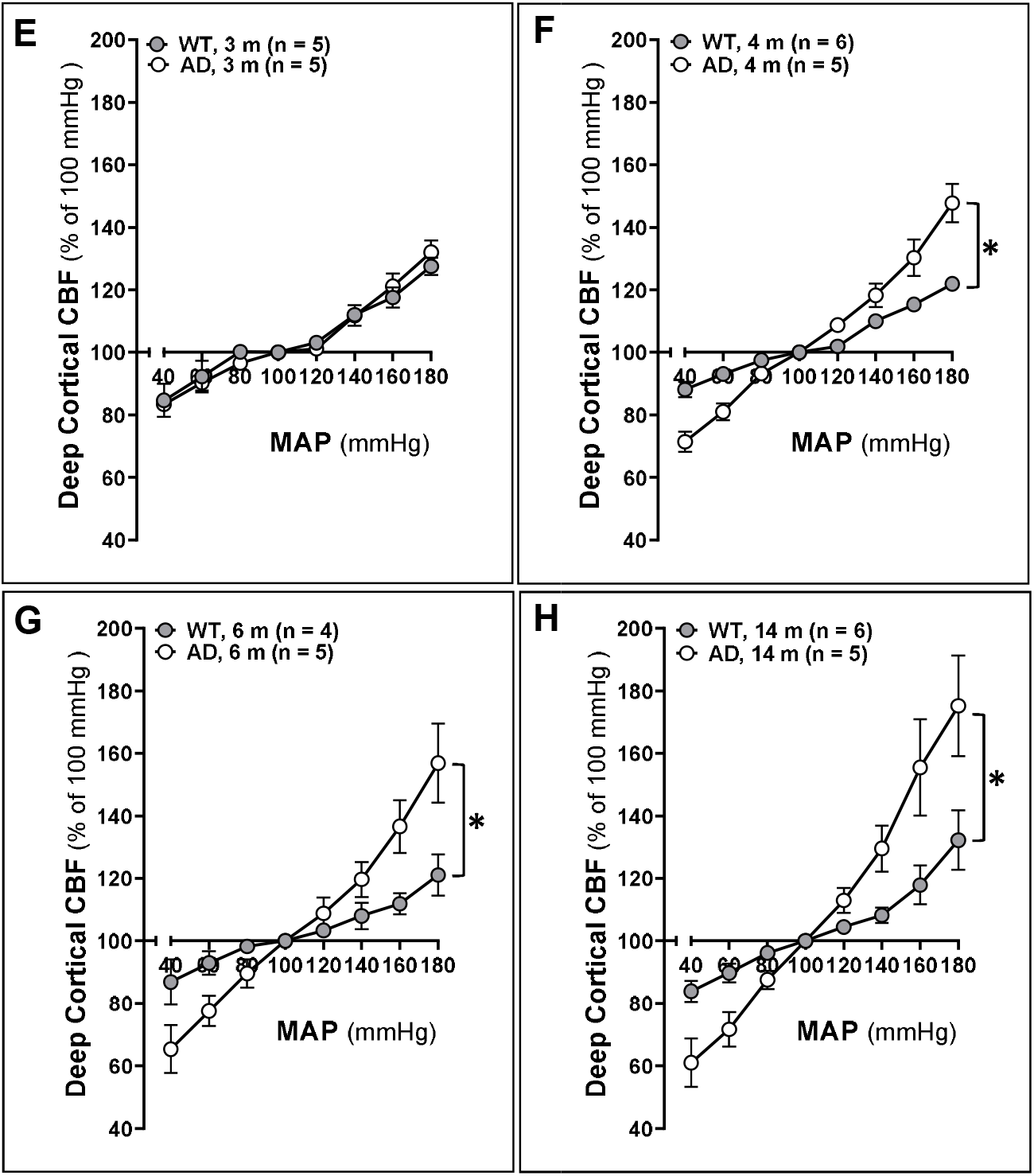
Cerebral blood flow (CBF) autoregulation. Comparison of the CBF autoregulation in 3, 4, 6, and 14 months old wildtype (WT; F344) and TgF344-AD (AD) rats. **A-D**. Comparison of surface cortical CBF autoregulation as of % to 100 mmHg. **E-H**. Comparison of deep cortical CBF autoregulation as of % to 100 mmHg. Data are presented as mean values ± SEM. Numbers in parentheses indicate the number of rats studied per group. * indicates a significant difference (*P* < 0.05) from the corresponding values in age-matched WT rats.

Deep cortical CBF autoregulation in 3, 4, 6, and 14 months male AD and WT rats was also compared (**Fig. 3E-H**). Similar to what we observed in surface cortical CBF autoregulation, WT and AD rats exhibited an intact deep cortical CBF autoregulation at 3 months of age. AD rats started to fail to autoregulate at 4 months, as CBF increased by 15.30 ± 1.78 % (WT) vs. 30.26 ± 5.84 % (AD) when perfusion pressure increased from 100 to 160 mmHg and decreased by 6.91± 1.99 % (WT) vs. 18.99 ± 2.67 % (AD) when perfusion pressure reduced from 100 to 60 mmHg, respectively. The impaired deep cortical CBF autoregulation was exacerbated with age in AD rats. AD rats exhibited autoregulatory breakthroughs above 140 mmHg at 4 months and above 120 mmHg at 6 and 14 months of age, respectively.

Results of CBF autoregulation obtained by LDF were confirmed using an LSI system. As presented in **Fig. 4**, brain perfusion was similar at pressure 60, 100, and 160 mmHg in 3 months old WT and AD rats when these rats exhibited intact CBF autoregulation. Hypoperfusion was observed around and below physiological pressures at 100 and 60 mmHg, and excessive CBF was detected at 160 mmHg (above autoregulatory breakthrough point) in 14 months AD rats, implying BBB breakdown.

**Figure 4.**
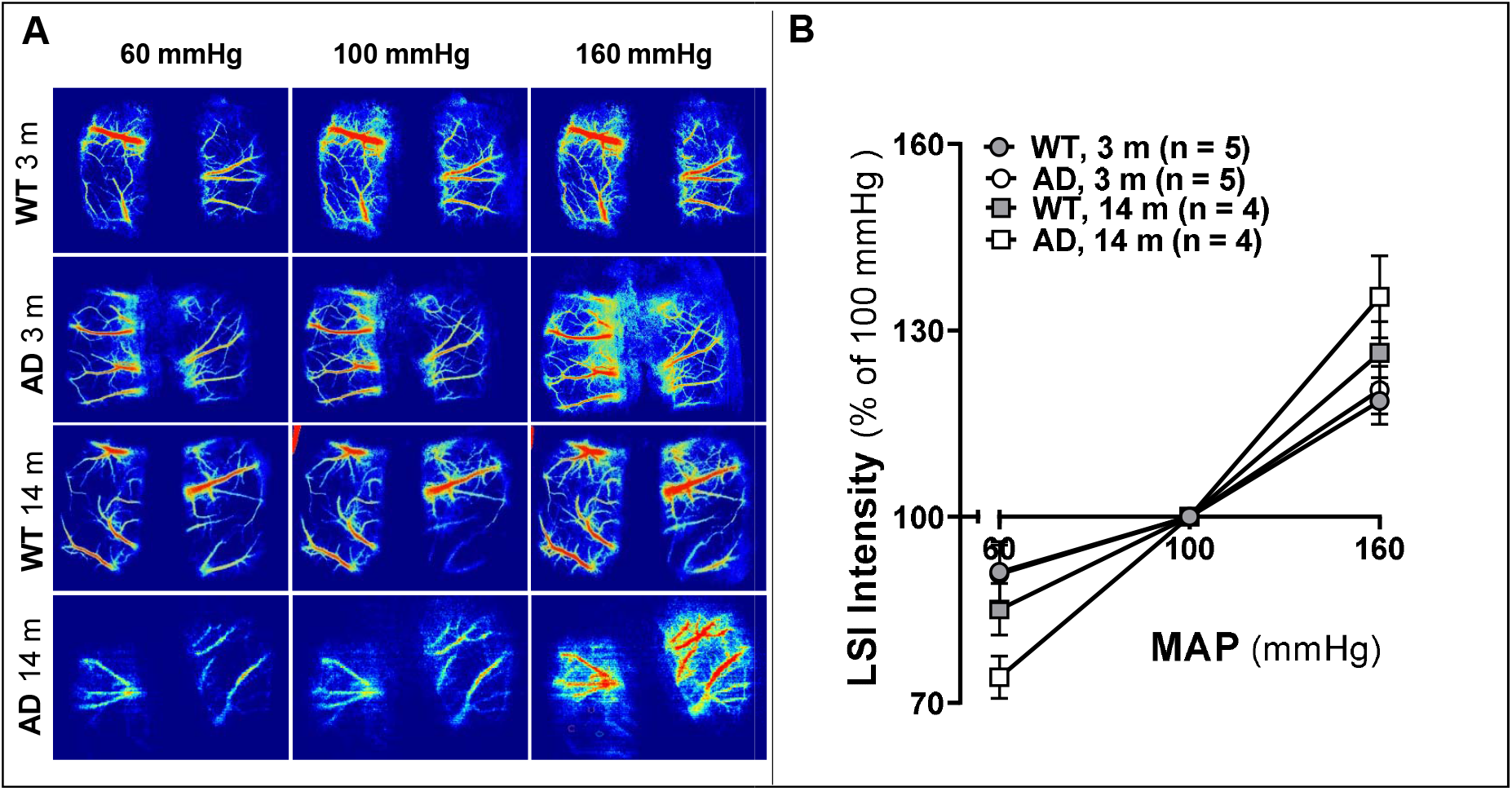
Brain Perfusion. Comparison of surface brain perfusion in 3 and 14 months old wildtype (WT; F344) and TgF344-AD (AD) rats using laser speckle imaging (LSI) system at pressure 60, 100, and 160 mmHg. **A**. Representatively laser speckle contrast images of surface cortical cerebral blood flow (CBF). **B**. Quantitation of LSI intensity as of % to 100 mmHg as an indicator of surface cortical CBF. Data are presented as mean values ± SEM. Numbers in parentheses indicate the number of rats studied per group. * indicates a significant difference (*P* < 0.05) from the corresponding values in age-matched WT rats.

### VSMC Contraction Assay

Changes in cerebral VSMC functional contractility could affect the MR of cerebral vasculature and CBF autoregulation associated with AD/ADRD ^57,61,62^. We compared the contractile capability of VSMCs isolated from MCAs of AD and WT rats. AD cerebral VSMCs displayed a weaker contraction index than WT, and the gel size was reduced by 3.56 ± 0.33 % and 8.32 ± 0.42 %, respectively (**Fig. 5**). We also compared cell contractility of WT cerebral VSMC treated with vehicle and Aβ (1-42). We first determined the time and doses of Aβ (1-42) treatments (data not shown). We then found that 24-hour administration of Aβ (1-42) reduced cell contraction of WT cerebral VSMCs by 5.84 ± 0.64 % (0.1 μM) and 5.98 ± 0.47 % (1 μM).

**Figure 5.**
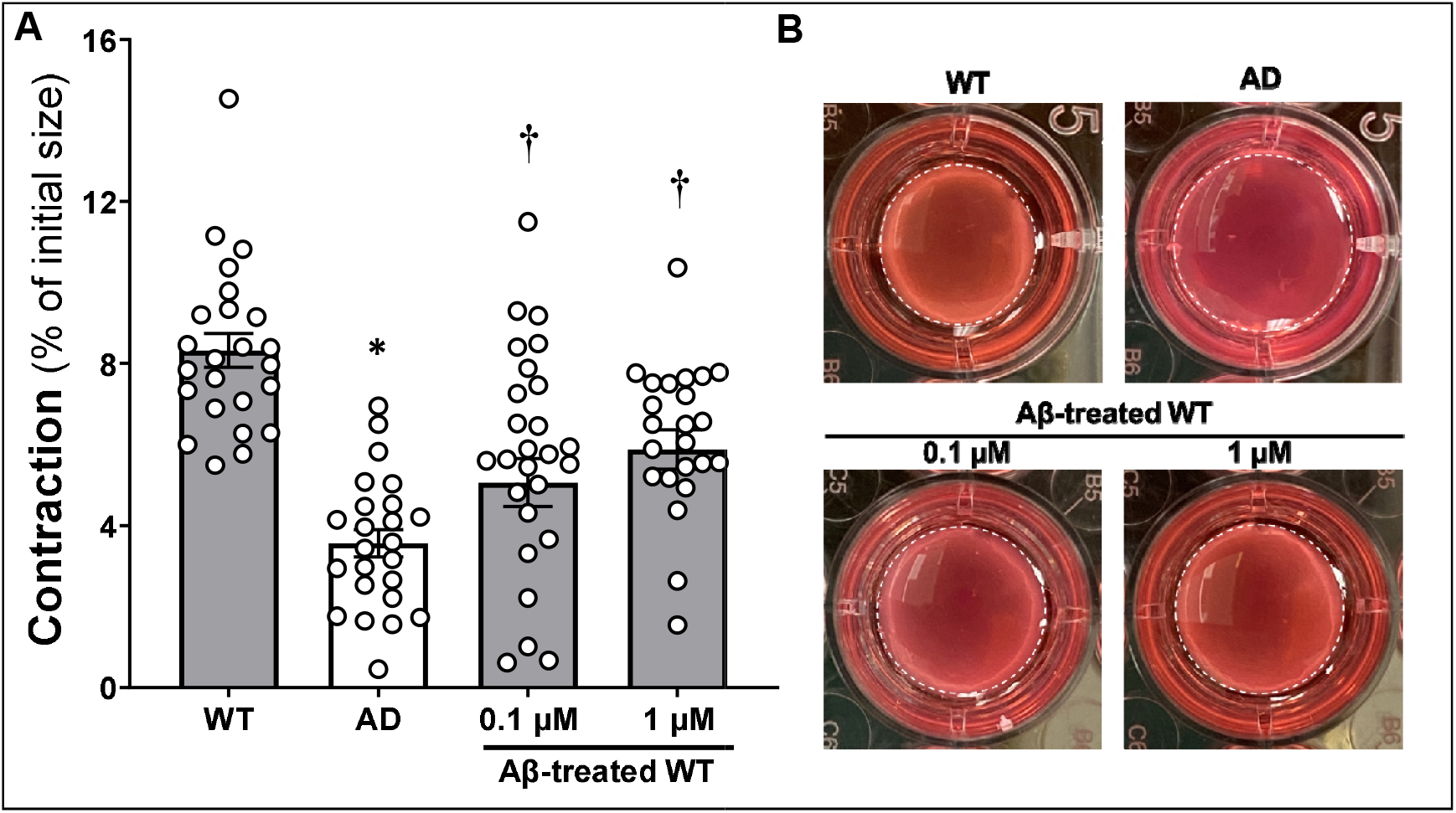
Cell Contractile Capability. Comparison of contractile capability in vascular smooth muscle cells (VSMCs) isolated from the middle cerebral artery of wildtype (WT; F344) and TgF344-AD (AD) rats, as well as the effect of Aβ (1-42) on the contractile capability of WT cerebral VSMCs. **A**. Quantitation of % constriction relative to the initial gel size of cerebral VSMCs in various groups and treatments. **B**. Representative images. Cerebral VSMCs were isolated from 3-4 rats per strain. Experiments using primary cerebral VSMCs were performed in triplicate and repeated three times. Mean values ± SEM are presented. Each dot (open circle) represents a single value obtained from an individual animal. * indicates *P* < 0.05 from the corresponding values in WT cells. **†** indicates *P* <0.05 from the corresponding values in WT cells without Aβ (1-42) treatments.

### Production of ROS and Mitochondrial Superoxide

As presented in **Fig. 6A and 6B**, DHE fluorescence intensity and DHE positive area per cell were 0.28- and 0.24-fold higher in VSMCs isolated from AD than WT rats, respectively. Similarly, MitoSOX fluorescence intensity and MitoSOX positive area per cell were 0.25- and 0.30-fold higher in VSMCs isolated from AD than WT rats, respectively (**Fig. 6C and 6D**). These results demonstrated that the production of ROS and mitochondrial superoxide was enhanced in AD cerebral VSMCs, which may contribute to impaired MR and CBF autoregulation ^9,57^.

**Figure 6.**
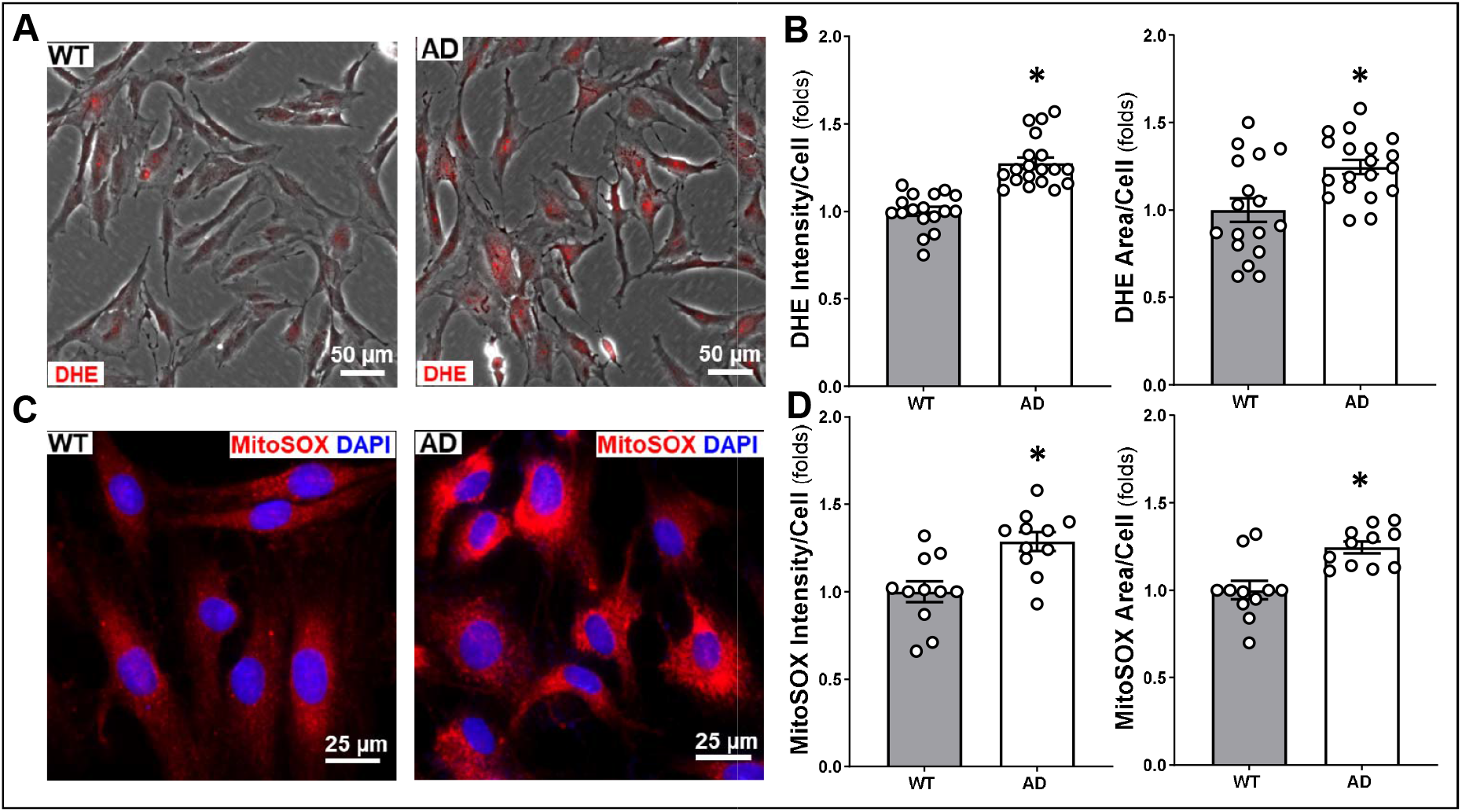
Production of Reactive Oxygen Species (ROS) and Mitochondrial Superoxide. The production of ROS and mitochondrial superoxide in vascular smooth muscle cells (VSMCs) isolated from the middle cerebral artery of wildtype (WT; F344) and TgF344-AD (AD) rats were compared. **A**. Representative images of dihydroethidium (DHE) staining as an indicator of ROS production. **B**. Quantitation of DHE fluorescence intensity and DHE positive area per cell. **C**. Representative images of MitoSOX™ as an indicator of mitochondrial superoxide production. **D**. Quantitation of MitoSOX™ fluorescence intensity and MitoSOX™ positive area per cell. Cerebral VSMCs were isolated from 3-4 rats per strain. Experiments using primary cerebral VSMCs were performed in triplicate and repeated three times. Mean values ± SEM are presented. Each dot (open circle) represents a single value obtained from an individual well. * indicates *P* < 0.05 from the corresponding values in WT cells.

### Mitochondrial Respiration and ATP Production

Changes in mitochondrial respiration and ATP production in cerebral VSMCs isolated from AD vs. WT rats were evaluated using a Seahorse XFe^24^ Extracellular Flux Analyzer by comparing OCR following our optimized protocols ^57,58^. As presented in **Fig. 7**, mitochondrial basal respiration (3,556.00 ± 164.37 vs. 4,171.67 ± 97.45 pmol/min/mg) and ATP production (2,988.77 ± 161.50 vs. 3,433.80 ± 56.52 pmol/min/mg) were significantly reduced in AD compared to WT cerebral VSMCs. Additionally, AD cerebral VSMCs presented significantly lower maximal respiration and lowered spare respiratory capacity than WT cells.

**Figure 7.**
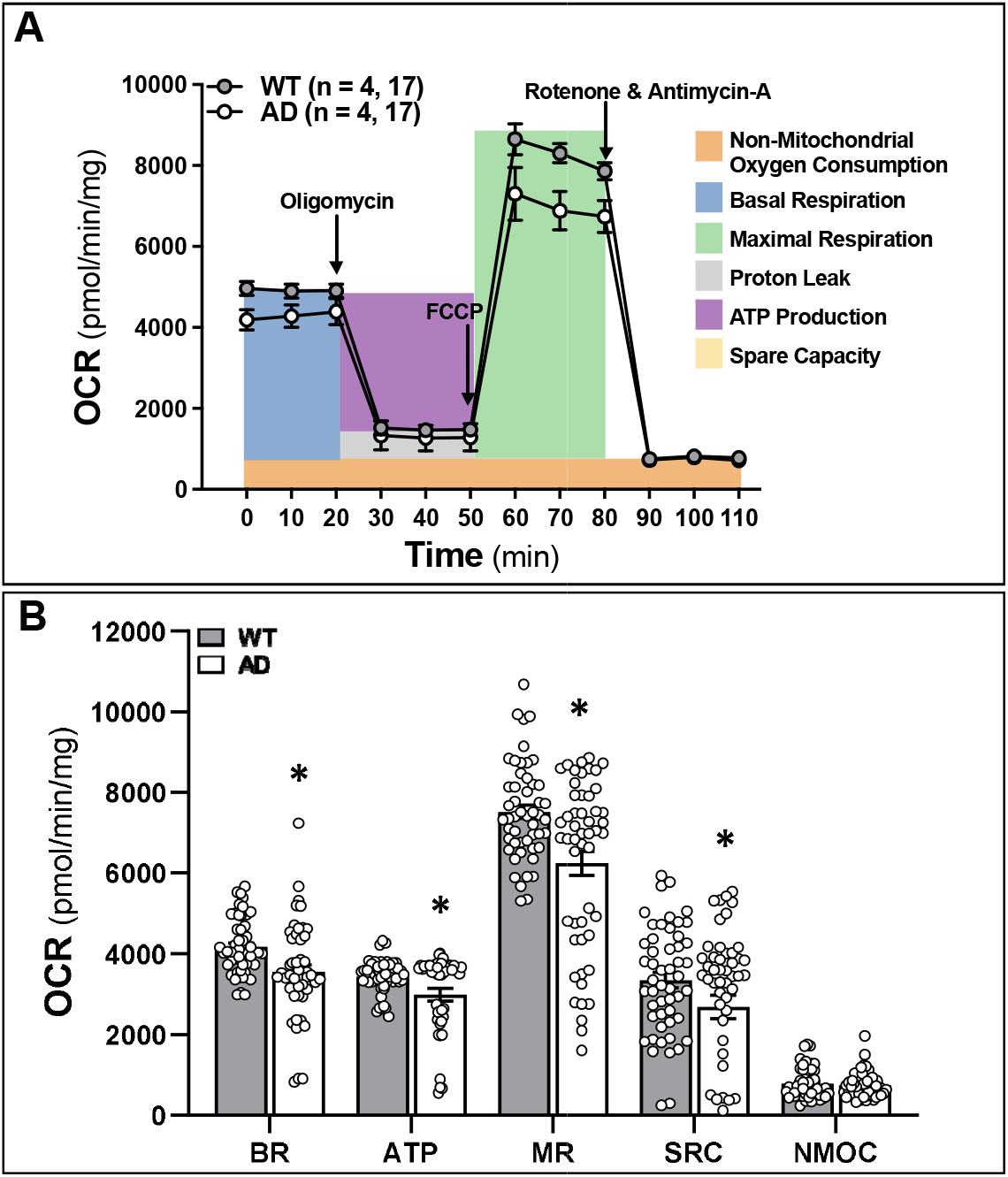
Mitochondrial Respiration and ATP Production. Mitochondrial respiration and ATP production in vascular smooth muscle cells (VSMCs) isolated from the middle cerebral artery of wildtype (WT; F344) and TgF344-AD (AD) rats were compared using the Seahorse XFe24 Extracellular Flux Analyzer by comparing the oxygen consumption rate (OCR). **A**. OCR Mito Stress test Seahorse profiles. Numbers in parentheses indicate the number of rats studied per group. **B**. Quantitative analysis of OCR. Each dot (open circle) represents a single value obtained from an individual well. Cerebral VSMCs were isolated from 3-4 rats per strain. Experiments using primary cerebral VSMCs were performed in triplicate and repeated three times. Mean values ± SEM are presented. * indicates *P* < 0.05 from the corresponding values in WT cells. BR, basal respiration; ATP, adenosine triphosphate; MR, maximal respiration; SRC, spare respiratory capacity; NMOC, non-mitochondrial oxygen consumption.

### Contractile Units in AD and WT Cerebral VSMCs

Disruption of the stabilization of the actin cytoskeleton by altering the F/G-actin ratio diminishes MLC phosphorylation and has been reported to reduce cell constriction in VSMCs and other motile cells ^36,39,63-66^. We next compared the structure, expression, and intracellular distribution of F-actin and MLC in AD and WT cerebral VSMCs. As shown in **Fig. 8**, the actin cytoskeleton displayed irregular polygonal shapes in the cell bodies of AD cerebral VSMCs compared with regular parallel stress fibers seen in WT cells. The expression of MLC, determined by fluorescence intensity, was lower in AD VSMCs. The MLC/F-actin ratio, indicating the numbers of the actin-myosin contractile units, was also reduced in AD cells.

**Figure 8.**
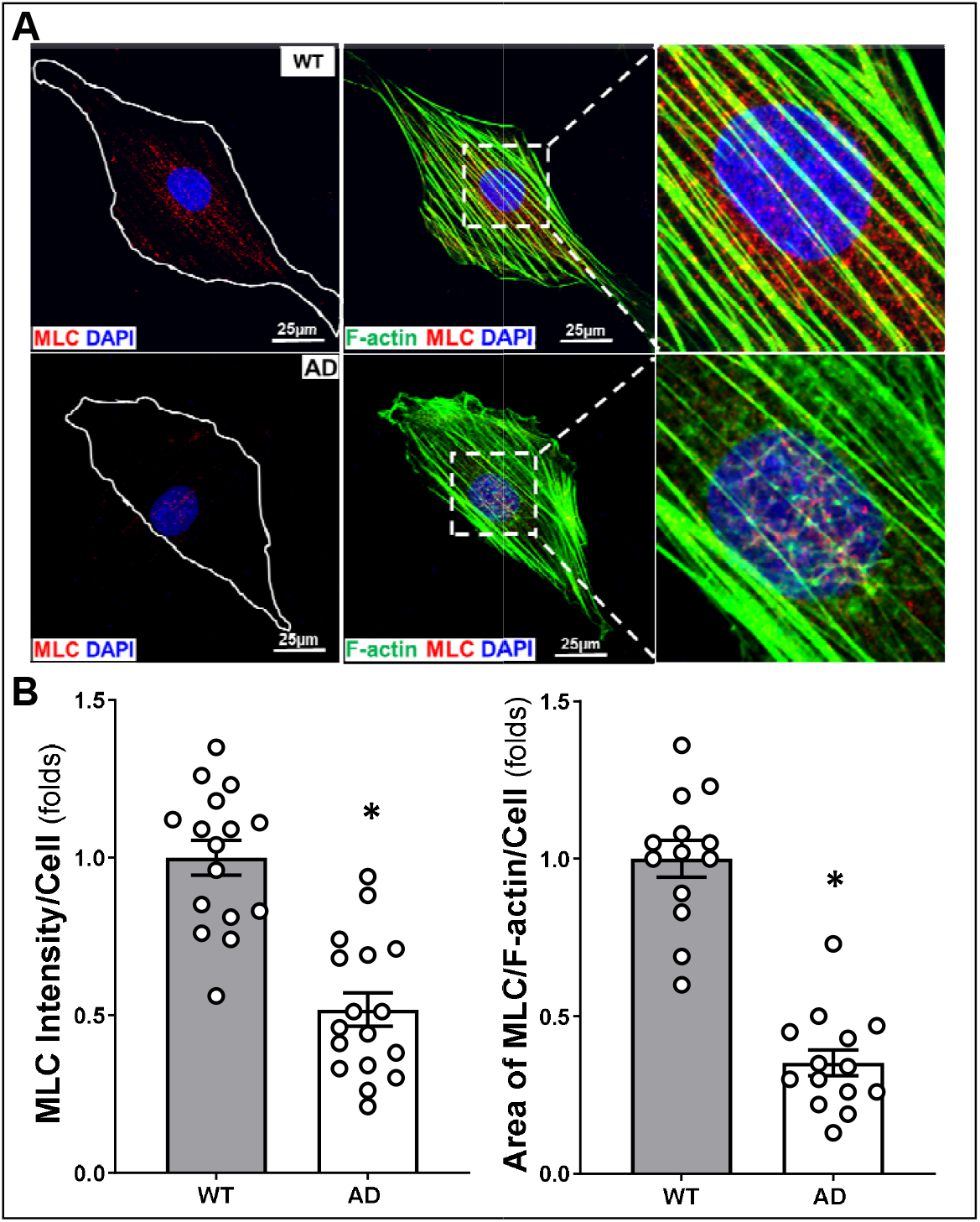
Contractile Units. The structure, expression, and intracellular distribution of F-actin and myosin light chain (MLC) in vascular smooth muscle cells (VSMCs) isolated from the middle cerebral artery of wildtype (WT; F344) and TgF344-AD (AD) rats were compared by immunocytochemistry. **A**. Representative images of immunostaining using Alexa Fluor 488 phalloidin for F-actin and the antibody against MLC. The cell nucleus was stained with 4′,6-diamidino-2-phenylindole (DAPI). **B**. Quantitative analysis of MLC fluorescence intensity and the MLC/F-actin ratios (of positively stained cells) as indicators of MLC expression and the numbers of the actin-myosin contractile units, respectively. Each dot (open circle) represents a single value obtained from an individual well. Cerebral VSMCs were isolated from 3-4 rats per strain. Experiments using primary cerebral VSMCs were performed in triplicate and repeated three times. Mean values ± SEM are presented. * indicates *P* < 0.05 from the corresponding values in WT cells.

## DISCUSSION

AD/ADRD is the most costly disease and one of the major causes of disability and morbidity in the elderly worldwide^1^. Although Aβ and Tau abnormality, neuro-inflammation, and oxidative stress have been considered to be the driving forces of AD/ADRD, more than 50% of AD/ADRD patients have mixed brain pathologies ^3,16^. Cholinesterase inhibitors, approved by the U.S. Food and Drug Administration (FDA) for AD treatments for AD, only alleviate AD symptoms by temporarily improving memory and cognition. Clinical trials targeting to correct Aβ and Tau abnormalities have mixed results ^67^. Aducanumab (Aduhelm^®^) was approved by the FDA in 2021 and has been reported to be able to reduce Aβ aggregation and improve cognition ^68^. More recently, results from a double-blind phase 3 clinical trial using another monoclonal antibody, Lecanemab, demonstrated that this drug that targets soluble Aβ soluble protofibrils could reduce cognitive decline in the early stages of AD ^69^. This evidence suggested that targeting to reduce Aβ accumulation could delay or reverse neuronal damage and dementia and thus could be a potential AD treatment. On the other hand, numerous studies also have indicated that neuronal damage and dementia in AD/ADRD are linked to brain hypoperfusion and cerebral vascular dysfunction ^4,5,70^. However, the chicken-or-egg relationship between Aβ accumulation and brain hypoperfusion has yet to be fully understood ^15,16^.

In the present study, we utilized the TgF344-AD rat and longitudinally characterized changes in cerebral hemodynamics in male AD rats at 3, 4, 6, and 14 months. This design is based on previous reports that TgF344-AD rats started to develop Aβ plaque, gliosis, and learning dysfunction at 6 months of age despite only containing mutant Aβ-producing genes ^17,18,71^. Of note, since there are sex differences in cerebral vascular structure and function and cognition ^40,72,73^, thus, in this study, we only focused on males to obtain a longitudinal characterization of the MR and autoregulation in the cerebral circulation in AD vs. WT rats. We first confirmed that AD rats developed cognitive deficits at 6 months without alteration of any other major biophysical parameters. We found that body weights, levels of plasma glucose and HbA1c, and MAP were similar in AD and age-matched WT rats across all ages. These results not only validated the reported phenotypes in this AD model ^17,22^, but also demonstrated that the cerebral vascular and cognitive dysfunction in this AD rat model was likely due to overexpression of Aβ in the absence of AD cardiovascular risk factors, such as hypertension, diabetes, and obesity ^30,31,74,75^.

It has been reported that cerebral vascular dysfunction often precedes the accumulation of Aβ and tau and cognitive decline in AD patients and animal models ^15,16^. Brain hypoperfusion could be attributed to impaired cerebral hemodynamics, BBB leakage, and NVU dysfunction.^4,61^ We found that the MR of the MCA and PA was intact in AD rats at 3 months of age, similar to what was seen in WT rats. AD rats started to lose the vasoconstrictive response to pressures at 4 months in both the MCA and PA. The impaired MR of MCA and PA was exacerbated with age in AD rats. AD vessels exhibited forced dilation over a pressure range of 120-180 mmHg (MCA) or 40-60 mmHg (PA) at 4 and 6 months of age and 100-180 mmHg (MCA) or 30-60 mmHg (PA) at 14 months of age. Results from our *in vivo* CBF autoregulation study were similar to the *ex vivo* MR study. We found that WT and AD rats exhibited a similar intact surface and deep cortical CBF autoregulation at 3 months of age. However, AD rats started to fail to autoregulate CBF at 4 months of age. The impaired surface and deep cortical CBF autoregulation were exacerbated with age in AD rats. AD surface cortical vessels exhibited autoregulatory breakthroughs above 140 and 120 mmHg at 4, and 6, and 14 months of age, respectively. AD deep cortical vessels exhibited autoregulatory breakthroughs above 140 mmHg at 4 months and above 120 mmHg at 6 and 14 months of age, respectively. These results indicated that impaired MR and CBF autoregulation appeared at least two months prior to the onset of cognitive deficits in the TgF344-AD rat. Our results also demonstrate that the MR and CBF autoregulation were impaired to a greater extent in old vs. young AD rats and in the deep vs. surface cortex, similar to what we found in diabetic rats ^9^. These results were further validated with brain perfusion using LSI. We found that 3 months old AD rats had similar brain perfusion to WT rats when perfusion pressure was at 60, 100, and 160 mmHg. Brain hypoperfusion was detected when perfusion pressure was at 100 and 60 mmHg, and BBB leakage was suggested when perfusion pressure was above the autoregulatory breakthrough point in 14 months AD rats.

We previously reported that old diabetic rats also exhibited impaired cerebral hemodynamics and cognitive impairments ^9^. We have evidence that the impaired cerebral hemodynamics in diabetic rats results from VSMC and arteriolar pericyte contractile dysfunction, possibly due to enhanced oxidative stress, mitochondrial dysfunction, and ATP depletion ^9,57,58^. We found abolished cell contractility in VSMCs isolated from AD MCA compared to cells obtained from WT vessels. Aβ (1-42) at two concentrations (0.1 and 1 μM) reduced cell contraction of WT cells. ROS production and mitochondrial ROS were enhanced, and mitochondrial respiration and ATP production were reduced in AD VSMCs compared with WT cells. Furthermore, we found that the stabilization of the actin cytoskeleton was disrupted, and the expression of MLC and the numbers of the actin-myosin contractile units were reduced in cerebral VSMCs isolated from AD rats. These results suggest that impaired cerebral hemodynamics in AD rats attributed to reduced cerebral VSMC functional contractility, which was associated with enhanced ROS production and reduced mitochondrial respiration and ATP production, similar to what we reported in diabetic-related dementia.

Other studies using AD animal models of AD suggested that Aβ may decrease the MR, CBF, and vasodilator responses ^76-78^. Our results of Aβ (1-42) reduced VSMC contractility indicate that it may have a direct effect on lowering cerebral vasoconstriction, consistent with previous findings ^79^. Surprisingly, Dietrich et al. ^80^ reported that Aβ-treated PAs of SD rats and in PAs of Tg2576 mice exhibited diminished dilation to ATP. They also found that both Aβ (1-40) and Aβ (1-42) constricted PAs isolated from Sprague-Dawley (SD) rats associated with increased ROS production. The Attwell group reported that Aβ evoked capillary pericyte-mediated vasoconstriction ^81,82^. PAs have been reported to exhibit a greater myogenic tone compared to cerebral arteries due to the lack of BK channels and enhanced L-type voltage-dependent calcium channel activity ^50^. The PA also acts as the “bottleneck” in the deep cortex that reduces CBF in the capillaries when constricted ^83^. Since the contractile cells on the wall of the MCA, PA, and capillary are VSMCs, mixed VSMCs and pericytes, and pericytes, respectively, it is possible that Aβ causes diverse effects, such as dilating VSMCs and constricting pericytes, along the cerebral vascular beds. Nevertheless, the effects of Aβ on the MCA, PA, and capillary may, directly or indirectly (via oxygen radicals), contribute to CBF reduction that plays a crucial role in driving cognitive impairments. On the other hand, brain hypoperfusion caused by Aβ may also evoke or amplify the production and/or reduce the clearance of Aβ. However, uncovering the underlying mechanisms need further study.

In conclusion, the present study underscores longitudinal changes in cerebral hemodynamics in the TgF344-AD transgenic rats and establishes that cerebral vascular dysfunction precedes learning and memory disturbance. The major findings of this study indicate that the TgF344-AD rats displayed significantly impaired MR and CBF autoregulation two months prior to learning and memory deficits; the dysfunction of cerebral hemodynamics in AD is exacerbated with age associated with reduced cerebral perfusion; abolished cell contractility contributes to cerebral hemodynamics imbalance in AD, which may be attributed to enhanced ROS production, decreased mitochondrial respiration and ATP depletion, and disrupted actin cytoskeleton in cerebral VSMCs. These findings highlight the complex and multifactorial nature of AD pathology, supporting the development of therapeutic interventions targeted at restoring cerebrovascular function in the early stage of AD.

## AUTHOR CONTRIBUTIONS

XF and FF conceived and designed research; XF, HZ, JB, YL, SMS, and YL performed experiments; XF, HZ, CT, JB, YL, and FF analyzed data; XF, CT, RJR, and FF interpreted results of experiments; XF, CT, and FF prepared figures; XF and FF drafted the manuscript; CT, SMS, HY, RJR, and FF edited and revised manuscript; all authors approved the final version of the manuscript.

## REFERENCES

1. Gaugler J, James, B., Johnson, T., Reimer, J., Solis, M., Weuve, J. 2022 Alzheimer’s disease facts and figures. Alzheimer’s & Dementia. 2022;18:700–789. doi: https://doi.org/10.1002/alz.12638

2. Sweeney MD, Kisler K, Montagne A, Toga AW, Zlokovic BV. The role of brain vasculature in neurodegenerative disorders. Nat Neurosci. 2018;21:1318–1331. doi: 10.1038/s41593-018-0234-x

3. Fang X, Crumpler RF, Thomas KN, Mazique JN, Roman RJ, Fan F. Contribution of cerebral microvascular mechanisms to age-related cognitive impairment and dementia. Physiol Int. 2022. doi: 10.1556/2060.2022.00020

4. Fan F, Roman RJ. Reversal of cerebral hypoperfusion: a novel therapeutic target for the treatment of AD/ADRD? Geroscience. 2021;43:1065–1067. doi: 10.1007/s11357-021-00357-7

5. Bracko O, Cruz Hernández JC, Park L, Nishimura N, Schaffer CB. Causes and consequences of baseline cerebral blood flow reductions in Alzheimer’s disease. J Cereb Blood Flow Metab. 2021;41:1501–1516. doi: 10.1177/0271678x20982383

6. Ungvari Z, Tarantini S, Donato AJ, Galvan V, Csiszar A. Mechanisms of Vascular Aging. Circ Res. 2018;123:849–867. doi: 10.1161/CIRCRESAHA.118.311378

7. Zhang H, Roman R, Fan F. Hippocampus is more susceptible to hypoxic injury: has the Rosetta Stone of regional variation in neurovascular coupling been deciphered? GeroScience. 2021. doi: 10.1007/s11357-021-00449-4

8. Fan F, Booz GW, Roman RJ. Aging diabetes, deconstructing the cerebrovascular wall. Aging (Albany NY). 2021;13:9158–9159. doi: 10.18632/aging.202963

9. Wang S, Lv W, Zhang H, Liu Y, Li L, Jefferson JR, Guo Y, Li M, Gao W, Fang X, et al. Aging exacerbates impairments of cerebral blood flow autoregulation and cognition in diabetic rats. Geroscience. 2020;42:1387–1410. doi: 10.1007/s11357-020-00233-w

10. Fan F, Geurts AM, Murphy SR, Pabbidi MR, Jacob HJ, Roman RJ. Impaired myogenic response and autoregulation of cerebral blood flow is rescued in CYP4A1 transgenic Dahl salt-sensitive rat. Am J Physiol Regul Integr Comp Physiol. 2015;308:R379–390. doi: 10.1152/ajpregu.00256.2014

11. Fan F, Pabbidi M, Lin RCS, Ge Y, Gomez-Sanchez EP, Rajkowska GK, Moulana M, Gonzalez-fernandez E, Sims J, Elliott MR, et al. Impaired myogenic response of MCA elevates transmission of pressure to penetrating arterioles and contributes to cerebral vascular disease in aging hypertensive FHH rats. The FASEB Journal. 2016;30:953.957-953.957. doi: https://doi.org/10.1096/fasebj.30.1_supplement.953.7

12. Gonzalez-Fernandez E, Liu Y, Auchus Alexander P, Fan F, Roman R. Vascular contributions to cognitive impairment and dementia: the emerging role of 20-HETE. Clinical Science. 2021;135:1929–1944. doi: 10.1042/CS20201033

13. Gonzalez-Fernandez E, Staursky D, Lucas K, Nguyen BV, Li M, Liu Y, Washington C, Coolen LM, Fan F, Roman RJ. 20-HETE Enzymes and Receptors in the Neurovascular Unit: Implications in Cerebrovascular Disease. Front Neurol. 2020;11:983. doi: 10.3389/fneur.2020.00983

14. Gonzalez-Fernandez E, Fan L, Wang S, Liu Y, Gao W, Thomas KN, Fan F, Roman RJ. The adducin saga: Pleiotropic genomic targets for precision medicine in human hypertension; vascular, renal, and cognitive diseases. Physiol Genomics. 2021. doi: 10.1152/physiolgenomics.00119.2021

15. Wang S, Mims PN, Roman RJ, Fan F. Is Beta-Amyloid Accumulation a Cause or Consequence of Alzheimer’s Disease? J Alzheimers Parkinsonism Dement. 2016;1.

16. Fang X, Zhang J, Roman RJ, Fan F. From 1901 to 2022, how far are we from truly understanding the pathogenesis of age-related dementia? GeroScience. 2022;44:1879–1883. doi: 10.1007/s11357-022-00591-7

17. Cohen RM, Rezai-Zadeh K, Weitz TM, Rentsendorj A, Gate D, Spivak I, Bholat Y, Vasilevko V, Glabe CG, Breunig JJ, et al. A transgenic Alzheimer rat with plaques, tau pathology, behavioral impairment, oligomeric aβ, and frank neuronal loss. J Neurosci. 2013;33:6245–6256. doi: 10.1523/jneurosci.3672-12.2013

18. Sare RM, Cooke SK, Krych L, Zerfas PM, Cohen RM, Smith CB. Behavioral Phenotype in the TgF344-AD Rat Model of Alzheimer’s Disease. Front Neurosci. 2020;14:601. doi: 10.3389/fnins.2020.00601

19. Dickie BR, Boutin H, Parker GJM, Parkes LM. Alzheimer’s disease pathology is associated with earlier alterations to blood-brain barrier water permeability compared with healthy ageing in TgF344-AD rats. NMR Biomed. 2021;34:e4510. doi: 10.1002/nbm.4510

20. Joo IL, Lai AY, Bazzigaluppi P, Koletar MM, Dorr A, Brown ME, Thomason LA, Sled JG, McLaurin J, Stefanovic B. Early neurovascular dysfunction in a transgenic rat model of Alzheimer’s disease. Sci Rep. 2017;7:46427. doi: 10.1038/srep46427

21. Chaney AM, Lopez-Picon FR, Serrière S, Wang R, Bochicchio D, Webb SD, Vandesquille M, Harte MK, Georgiadou C, Lawrence C, et al. Prodromal neuroinflammatory, cholinergic and metabolite dysfunction detected by PET and MRS in the TgF344-AD transgenic rat model of AD: a collaborative multi-modal study. Theranostics. 2021;11:6644–6667. doi: 10.7150/thno.56059

22. Bazzigaluppi P, Beckett TL, Koletar MM, Hill ME, Lai A, Trivedi A, Thomason L, Dorr A, Gallagher D, Librach CL, et al. Combinatorial Treatment Using Umbilical Cord Perivascular Cells and Aβ Clearance Rescues Vascular Function Following Transient Hypertension in a Rat Model of Alzheimer Disease. Hypertension. 2019;74:1041–1051. doi: 10.1161/hypertensionaha.119.13187

23. Cipolla MJ. The Cerebral Circulation. San Rafael (CA): Morgan & Claypool Life Sciences. 2009. doi: https://www.ncbi.nlm.nih.gov/books/NBK53086/

24. Joo IL, Lam WW, Oakden W, Hill ME, Koletar MM, Morrone CD, Stanisz GJ, McLaurin J, Stefanovic B. Early alterations in brain glucose metabolism and vascular function in a transgenic rat model of Alzheimer’s disease. Prog Neurobiol. 2022;217:102327. doi: 10.1016/j.pneurobio.2022.102327

25. Ni A, Sethi A, Initiative ftAsDN. Functional Genetic Biomarkers of Alzheimer’s Disease and Gene Expression from Peripheral Blood. bioRxiv. 2021:2021.2001.2015.426891. doi: 10.1101/2021.01.15.426891

26. Yang AC, Vest RT, Kern F, Lee DP, Agam M, Maat CA, Losada PM, Chen MB, Schaum N, Khoury N, et al. A human brain vascular atlas reveals diverse mediators of Alzheimer’s risk. Nature. 2022. doi: 10.1038/s41586-021-04369-3

27. Fan F, Simino J, Auchus AP, Knopman DS, Boerwinkle E, Fornage M, Mosley TH, Roman RJ. Functional variants in CYP4A11 and CYP4F2 are associated with cognitive impairment and related dementia endophenotypes in the elderly. In: The 16th International Winter Eicosanoid Conference. Baltimore; 2016:CV5.

28. Thomas K, Wang S, Zhang H, Crumpler R, Elliott P, Ryu J, Fang X, Strong L, Liu Y, Zheng B, et al. Abstract 35: Gamma Adducin Dysfunction Leads To Cerebrovascular Distention, Blood Brain Barrier Leakage, And Cognitive Deficits In The Fawn-hooded Hypertensive Rats. Hypertension. 2021;78. doi: 10.1161/hyp.78.suppl_1.35

29. Claassen J, Thijssen DHJ, Panerai RB, Faraci FM. Regulation of Cerebral Blood Flow in Humans: Physiology and Clinical Implications of Autoregulation. Physiol Rev. 2021. doi: 10.1152/physrev.00022.2020

30. Shekhar S, Wang S, Mims PN, Gonzalez-Fernandez E, Zhang C, He X, Liu CY, Lv W, Wang Y, Huang J, et al. Impaired Cerebral Autoregulation-A Common Neurovascular Pathway in Diabetes may Play a Critical Role in Diabetes-Related Alzheimer’s Disease. Curr Res Diabetes Obes J. 2017;2:555587.

31. Shekhar S, Liu R, Travis OK, Roman RJ, Fan F. Cerebral Autoregulation in Hypertension and Ischemic Stroke: A Mini Review. J Pharm Sci Exp Pharmacol. 2017;2017:21–27.

32. Faraci FM, Heistad DD. Regulation of the cerebral circulation: role of endothelium and potassium channels. Physiol Rev. 1998;78:53–97. doi: 10.1152/physrev.1998.78.1.53

33. Harper SL, Bohlen HG, Rubin MJ. Arterial and microvascular contributions to cerebral cortical autoregulation in rats. Am J Physiol. 1984;246:H17–24. doi: 10.1152/ajpheart.1984.246.1.H17

34. Kontos HA, Wei EP, Navari RM, Levasseur JE, Rosenblum WI, Patterson JL, Jr. Responses of cerebral arteries and arterioles to acute hypotension and hypertension. Am J Physiol. 1978;234:H371–383. doi: 10.1152/ajpheart.1978.234.4.H371

35. Paulson OB, Strandgaard S, Edvinsson L. Cerebral autoregulation. Cerebrovascular and brain metabolism reviews. 1990;2:161–192.

36. Gao W, Liu Y, Fan L, Zheng B, Jefferson JR, Wang S, Zhang H, Fang X, Nguyen BV, Zhu T, et al. Role of γ-adducin in actin cytoskeleton rearrangements in podocyte pathophysiology. Am J Physiol Renal Physiol. 2021;320:F97–f113. doi: 10.1152/ajprenal.00423.2020

37. Wang S, Jiao F, Border JJ, Fang X, Crumpler RF, Liu Y, Zhang H, Jefferson J, Guo Y, Elliott PS, et al. Luseogliflozin, a sodium-glucose cotransporter-2 inhibitor, reverses cerebrovascular dysfunction and cognitive impairments in 18-mo-old diabetic animals. Am J Physiol Heart Circ Physiol. 2022;322:H246–H259. doi: 10.1152/ajpheart.00438.2021

38. Zhang C, He X, Murphy SR, Zhang H, Wang S, Ge Y, Gao W, Williams JM, Geurts AM, Roman RJ, et al. Knockout of Dual-Specificity Protein Phosphatase 5 Protects Against Hypertension-Induced Renal Injury. J Pharmacol Exp Ther. 2019;370:206–217. doi: 10.1124/jpet.119.258954

39. Fan F, Geurts AM, Pabbidi MR, Ge Y, Zhang C, Wang S, Liu Y, Gao W, Guo Y, Li L, et al. A Mutation in gamma-Adducin Impairs Autoregulation of Renal Blood Flow and Promotes the Development of Kidney Disease. J Am Soc Nephrol. 2020;31:687–700. doi: 10.1681/ASN.2019080784

40. Wang S, Zhang H, Liu Y, Li L, Guo Y, Jiao F, Fang X, Jefferson JR, Li M, Gao W, et al. Sex differences in the structure and function of rat middle cerebral arteries. Am J Physiol Heart Circ Physiol. 2020;318:H1219–H1232. doi: 10.1152/ajpheart.00722.2019

41. Fan F, Wang SX, Mims PN, Maeda KJ, Li LY, Geurts AM, Roman RJ. Knockout of matrix metalloproteinase-9 rescues the development of cognitive impairments in hypertensive Dahl salt sensitive rats. FASEB J. 2017;31:842.846-842.846. doi: 10.1096/fasebj.31.1_supplement.842.6

42. Wang SX, Jiao F, Guo Y, Zhang HW, He XC, Maranon RO, Alexandra B, Pabbidi M, Roman RJ, Fan F. Excessive salt consumption increases susceptibility to cerebrovascular dysfunction and cognitive impairments in the elderly of both sexes. FASEB J. 2019;33:511.517-511.517.

43. Bimonte HA, Hyde LA, Hoplight BJ, Denenberg VH. In two species, females exhibit superior working memory and inferior reference memory on the water radial-arm maze. Physiol Behav. 2000;70:311–317. doi: 10.1016/s0031-9384(00)00259-6

44. Hyde LA, Hoplight BJ, Denenberg VH. Water version of the radial-arm maze: learning in three inbred strains of mice. Brain Res. 1998;785:236–244. doi: 10.1016/s0006-8993(97)01417-0

45. Hyde LA, Sherman GF, Hoplight BJ, Denenberg VH. Working memory deficits in BXSB mice with neocortical ectopias. Physiol Behav. 2000;70:1–5. doi: 10.1016/s0031-9384(00)00239-0

46. Penley SC, Gaudet CM, Threlkeld SW. Use of an eight-arm radial water maze to assess working and reference memory following neonatal brain injury. J Vis Exp. 2013:50940. doi: 10.3791/50940

47. Nunez J. Morris Water Maze Experiment. J Vis Exp. 2008. doi: 10.3791/897

48. Zhang H, Zhang C, Liu Y, Gao W, Wang S, Fang X, Guo Y, Li M, Liu R, Roman RJ, et al. Influence of dual-specificity protein phosphatase 5 on mechanical properties of rat cerebral and renal arterioles. Physiol Rep. 2020;8:e14345. doi: 10.14814/phy2.14345

49. Cipolla MJ, Chan SL, Sweet J, Tavares MJ, Gokina N, Brayden JE. Postischemic reperfusion causes smooth muscle calcium sensitization and vasoconstriction of parenchymal arterioles. Stroke. 2014;45:2425–2430. doi: 10.1161/STROKEAHA.114.005888

50. Cipolla MJ, Sweet J, Chan SL, Tavares MJ, Gokina N, Brayden JE. Increased pressure-induced tone in rat parenchymal arterioles vs. middle cerebral arteries: role of ion channels and calcium sensitivity. J Appl Physiol (1985). 2014;117:53–59. doi: 10.1152/japplphysiol.00253.2014

51. Pires PW, Dabertrand F, Earley S. Isolation and Cannulation of Cerebral Parenchymal Arterioles. J Vis Exp. 2016. doi: 10.3791/53835

52. Fan F, Sun CW, Maier KG, Williams JM, Pabbidi MR, Didion SP, Falck JR, Zhuo J, Roman RJ. 20-Hydroxyeicosatetraenoic acid contributes to the inhibition of K+ channel activity and vasoconstrictor response to angiotensin II in rat renal microvessels. PLoS One. 2013;8:e82482. doi: 10.1371/journal.pone.0082482

53. Liu Y, Zhang H, Wu CY, Yu T, Fang X, Ryu JJ, Zheng B, Chen Z, Roman RJ, Fan F. 20-HETE-promoted cerebral blood flow autoregulation is associated with enhanced pericyte contractility. Prostaglandins Other Lipid Mediat. 2021;154:106548. doi: 10.1016/j.prostaglandins.2021.106548

54. Fan F, Geurts AM, Pabbidi MR, Smith SV, Harder DR, Jacob H, Roman RJ. Zinc-finger nuclease knockout of dual-specificity protein phosphatase-5 enhances the myogenic response and autoregulation of cerebral blood flow in FHH.1BN rats. PLoS One. 2014;9:e112878. doi: 10.1371/journal.pone.0112878

55. Korbo L, Pakkenberg B, Ladefoged O, Gundersen HJ, Arlien-Soborg P, Pakkenberg H. An efficient method for estimating the total number of neurons in rat brain cortex. Journal of neuroscience methods. 1990;31:93–100. doi: 10.1016/0165-0270(90)90153-7

56. He J, Lu H, Young L, Deng R, Callow D, Tong S, Jia X. Real-time quantitative monitoring of cerebral blood flow by laser speckle contrast imaging after cardiac arrest with targeted temperature management. J Cereb Blood Flow Metab. 2019;39:1161–1171. doi: 10.1177/0271678X17748787

57. Guo Y, Wang S, Liu Y, Fan L, Booz GW, Roman RJ, Chen Z, Fan F. Accelerated cerebral vascular injury in diabetes is associated with vascular smooth muscle cell dysfunction. Geroscience. 2020;42:547–561. doi: 10.1007/s11357-020-00179-z

58. Liu Y, Zhang H, Wang S, Guo Y, Fang X, Zheng B, Gao W, Yu H, Chen Z, Roman RJ, et al. Reduced pericyte and tight junction coverage in old diabetic rats are associated with hyperglycemia-induced cerebrovascular pericyte dysfunction. Am J Physiol Heart Circ Physiol. 2021;320:H549–H562. doi: 10.1152/ajpheart.00726.2020

59. Robinson KM, Janes MS, Beckman JS. The selective detection of mitochondrial superoxide by live cell imaging. Nat Protoc. 2008;3:941–947. doi: 10.1038/nprot.2008.56

60. Fan F, Pabbidi MR, Ge Y, Li L, Wang S, Mims PN, Roman RJ. Knockdown of Add3 impairs the myogenic response of renal afferent arterioles and middle cerebral arteries. Am J Physiol Renal Physiol. 2017;312:F971–F981. doi: 10.1152/ajprenal.00529.2016

61. Wang S, Tang C, Liu Y, Border JJ, Roman RJ, Fan F. Impact of impaired cerebral blood flow autoregulation on cognitive impairment. Frontiers in Aging. 2022;3. doi: 10.3389/fragi.2022.1077302

62. Fan F, Ge Y, Lv W, Elliott MR, Muroya Y, Hirata T, Booz GW, Roman RJ. Molecular mechanisms and cell signaling of 20-hydroxyeicosatetraenoic acid in vascular pathophysiology. Front Biosci (Landmark Ed). 2016;21:1427–1463.

63. Gunst SJ, Zhang W. Actin cytoskeletal dynamics in smooth muscle: a new paradigm for the regulation of smooth muscle contraction. Am J Physiol Cell Physiol. 2008;295:C576–587. doi: 10.1152/ajpcell.00253.2008

64. Noguchi TQ, Morimatsu M, Iwane AH, Yanagida T, Uyeda TQ. The role of structural dynamics of actin in class-specific myosin motility. PLoS One. 2015;10:e0126262. doi: 10.1371/journal.pone.0126262

65. Stricker J, Falzone T, Gardel ML. Mechanics of the F-actin cytoskeleton. J Biomech. 2010;43:9–14. doi: 10.1016/j.jbiomech.2009.09.003

66. Wang S, Jiao F, Guo Y, Booz G, Roman R, Fan F. Role of Vascular Smooth Muscle Cells in Diabetes-related Vascular Cognitive Impairment. Stroke. 2019;50:ATP556–ATP556.

67. Huang LK, Chao SP, Hu CJ. Clinical trials of new drugs for Alzheimer disease. J Biomed Sci. 2020;27:18. doi: 10.1186/s12929-019-0609-7

68. Tampi RR, Forester BP, Agronin M. Aducanumab: evidence from clinical trial data and controversies. Drugs Context. 2021;10. doi: 10.7573/dic.2021-7-3

69. van Dyck CH, Swanson CJ, Aisen P, Bateman RJ, Chen C, Gee M, Kanekiyo M, Li D, Reyderman L, Cohen S, et al. Lecanemab in Early Alzheimer’s Disease. N Engl J Med. 2022. doi: 10.1056/NEJMoa2212948

70. Crumpler R, Roman RJ, Fan F. Capillary Stalling: A Mechanism of Decreased Cerebral Blood Flow in AD/ADRD. J Exp Neurol. 2021;2:149–153. doi: 10.33696/neurol.2.048

71. Shekhar S, Travis OK, He X, Roman RJ, Fan F. Menopause and Ischemic Stroke: A Brief Review. MOJ Toxicol. 2017;3. doi: 10.15406/mojt.2017.03.00059

72. Upadhayay N, Guragain S. Comparison of cognitive functions between male and female medical students: a pilot study. J Clin Diagn Res. 2014;8:Bc12–15. doi: 10.7860/jcdr/2014/7490.4449

73. Jäncke L. Sex/gender differences in cognition, neurophysiology, and neuroanatomy. F1000Res. 2018;7. doi: 10.12688/f1000research.13917.1

74. Shekhar S, Varghese K, Li M, Fan L, Booz GW, Roman RJ, Fan F. Conflicting Roles of 20-HETE in Hypertension and Stroke. Int J Mol Sci. 2019;20:4500. doi: 10.3390/ijms20184500

75. Pegueroles J, Jiménez A, Vilaplana E, Montal V, Carmona-Iragui M, Pané A, Alcolea D, Videla L, Casajoana A, Clarimón J, et al. Obesity and Alzheimer’s disease, does the obesity paradox really exist? A magnetic resonance imaging study. Oncotarget. 2018;9:34691–34698. doi: 10.18632/oncotarget.26162

76. Christie R, Yamada M, Moskowitz M, Hyman B. Structural and functional disruption of vascular smooth muscle cells in a transgenic mouse model of amyloid angiopathy. Am J Pathol. 2001;158:1065–1071. doi: 10.1016/s0002-9440(10)64053-9

77. Zhang F, Eckman C, Younkin S, Hsiao KK, Iadecola C. Increased susceptibility to ischemic brain damage in transgenic mice overexpressing the amyloid precursor protein. J Neurosci. 1997;17:7655–7661. doi: 10.1523/jneurosci.17-20-07655.1997

78. Niwa K, Kazama K, Younkin L, Younkin SG, Carlson GA, Iadecola C. Cerebrovascular autoregulation is profoundly impaired in mice overexpressing amyloid precursor protein. Am J Physiol Heart Circ Physiol. 2002;283:H315–323. doi: 10.1152/ajpheart.00022.2002

79. Hald ES, Timm CD, Alford PW. Amyloid Beta Influences Vascular Smooth Muscle Contractility and Mechanoadaptation. J Biomech Eng. 2016;138. doi: 10.1115/1.4034560

80. Dietrich HH, Xiang C, Han BH, Zipfel GJ, Holtzman DM. Soluble amyloid-beta, effect on cerebral arteriolar regulation and vascular cells. Mol Neurodegener. 2010;5:15. doi: 10.1186/1750-1326-5-15

81. Korte N, Nortley R, Attwell D. Cerebral blood flow decrease as an early pathological mechanism in Alzheimer’s disease. Acta Neuropathologica. 2020;140:793–810. doi: 10.1007/s00401-020-02215-w

82. Nortley R, Korte N, Izquierdo P, Hirunpattarasilp C, Mishra A, Jaunmuktane Z, Kyrargyri V, Pfeiffer T, Khennouf L, Madry C, et al. Amyloid β oligomers constrict human capillaries in Alzheimer’s disease via signaling to pericytes. Science. 2019;365. doi: 10.1126/science.aav9518

83. Nishimura N, Schaffer CB, Friedman B, Lyden PD, Kleinfeld D. Penetrating arterioles are a bottleneck in the perfusion of neocortex. Proc Natl Acad Sci U S A. 2007;104:365–370. doi: 10.1073/pnas.0609551104

